# Enhancer-promoter interactions and transcription are maintained upon acute loss of CTCF, cohesin, WAPL, and YY1

**DOI:** 10.1101/2021.07.14.452365

**Authors:** Tsung-Han S. Hsieh, Claudia Cattoglio, Elena Slobodyanyuk, Anders S. Hansen, Xavier Darzacq, Robert Tjian

## Abstract

It remains unclear why acute depletion of CTCF and cohesin only marginally affects expression of most genes despite substantially perturbing 3D genome folding at the level of domains and structural loops. To address this conundrum, we used high-resolution Micro-C and nascent transcript profiling to find that enhancer-promoter (E-P) interactions are largely insensitive to acute (3-hour) depletion of CTCF, cohesin, and WAPL. YY1 has been proposed to be a structural regulator of E-P loops, but acute YY1 depletion also had minimal effects on E-P loops, transcription, and 3D genome folding. Strikingly, live-cell single-molecule imaging revealed that cohesin depletion reduced transcription factor binding to chromatin. Thus, although neither CTCF, cohesin, WAPL, nor YY1 are required for the short-term maintenance of most E-P interactions and gene expression, we propose that cohesin may serve as a “transcription factor binding platform” that facilitates transcription factor binding to chromatin.

## Introduction

High-throughput chromosomal conformation capture (Hi-C)-based assays have transformed our understanding of 3D genome folding^1,2^. Based on such studies, we can distinguish at least three levels of 3D genome folding. First, the genome is segregated into A and B compartmental domains, which largely correspond to active and inactive chromatin segments, respectively, and appear as a plaid-like pattern in Hi-C contact maps^3^. Second, the proteins CTCF and cohesin help fold the genome into topologically associating domains (TADs)^4,5^ and structural chromatin loops^6^. Third, at a much finer scale, transcriptional elements engage in long-range chromatin interactions such as enhancer-promoter (E-P) and promoter-promoter (P-P) interactions to form local domains^7–9^.

Elegant experiments combining acute protein depletion of CTCF, cohesin, and cohesin regulatory proteins with Hi-C or imaging approaches have revealed the role of CTCF and cohesin in regulating the first two levels, TADs and compartments^10–14^. These studies have shown that while CTCF and cohesin play only a minor role in compartmentalization, their removal largely eliminates TADs and chromatin loops anchored by these proteins across the genome. CTCF and cohesin are thought to form TADs and loops through loop extrusion^15,16^. This model posits that cohesin extrudes bidirectional loops until it encounters convergent and occupied CTCF binding sites. When averaged across cell populations, the extruded chromatin appears to be spatially organized into a self-interacting domain (TAD or loop domain), and CTCF binding sites constitute the domain boundaries that restrict inter-domain interactions. Halting of cohesin extrusion at CTCF sites sometimes gives rise to sharp corner peaks in contact maps, known as loops or corner dots. WAPL, a cohesin unloader, releases cohesin from chromatin and WAPL depletion therefore increases cohesin residence times as well as the amount of cohesin on chromatin^12,14^.

However, Hi-C is ineffective for capturing the third level of 3D genome folding, the fine-scale transcriptionally important E-P and P-P interactions ^7,17^. Indeed, a recent genetic screen found that the majority of functional E-P interactions are not identified as contacts in Hi-C data^18^. Our understanding of the role of CTCF and cohesin in regulating gene expression has thus mainly come from genetic experiments. Paradigmatic experiments focusing on mouse development suggested that TAD disruption through inversion or deletion of CTCF sites around developmental genes such as *Epha4*, *Kcnj*, or *Ihh* can cause severe limb malformation^19^. Similar studies at the locus encoding the morphogen Sonic hedgehog has led to somewhat inconsistent effects on transcription and developmental phenotypes, perhaps due to manipulation of different CTCF sites^20,21^. Furthermore, CTCF and cohesin appeared to be crucial to some biological processes such as neuronal maturation^22^ and lipopolysaccharide-induced inflammatory response^23^, but their presence seemed dispensable in other cases such as neuronal activity-dependent transcription^22^ and immune cell transdifferentiation^23^. Thus, it remained unclear if, when, where, and how CTCF/cohesin regulates E-P and P-P interactions and gene expression.

Our current lack of understanding of the role of CTCF, cohesin, and other factors in regulating E-P, P-P, and transcriptionally relevant fine-scale genome folding has largely been limited by the inability of Hi-C to directly interrogate these finer scale interactions. We recently reported that Micro-C can effectively resolve ultra-fine 3D genome folding at nucleosome resolution, including E-P and P-P interactions, thus overcoming this limitaion^7,24–27^. Here, we used Micro-C to systematically investigate fine-scale 3D genome folding before and after acute depletion (3 hours) of CTCF, the cohesin subunit RAD21, the cohesin unloader WAPL, and the putative structural protein YY1 in mouse embryonic stem cells (mESCs). We also profiled chromatin occupancy, total RNAs, and nascent transcripts^28^ in the same conditions. This integrated genome-wide fine-resolution mapping approach allowed us to dissect the primary effects of depleting loop extrusion factors on gene regulatory chromatin interactions and transcription. Finally, focusing on the dynamics of YY1 uncovered an unexpected role for cohesin in facilitating transcription factor (TF) binding.

## Results

### Genome-wide identification of transcription-linked chromatin loops

We previously used Micro-C to determine fine-scale (∼200 nucleotide) 3D genome structure at a resolution below TADs^7^. These structures correlate well with transcriptional activity and often facilitate contact between promoters and promoters (P-P) or enhancers and promoters (E-P), forming “dots” or “loops” at their intersections on Micro-C maps. Using a newly developed loop caller, Mustache^29^, we identified over 75,000 statistically significant dots/loops in mESCs, which is about 2.5 times greater than in our previous report^7^. Through analysis of local chromatin state enrichment at loop anchors, we sub-classified these loops into ∼13,735 cohesin loops, ∼20,369 E-P loops, ∼7,433 P-P loops, and ∼700 Polycomb-associated loops (**Fig. 1a-b and Extended Data Fig. 1a**). These chromatin loops span a broad range of lengths with a median size of ∼160 kb for cohesin loops and ∼100 kb for E-P or P-P loops (**Extended Data Fig. 1c**). Although cohesin loops exhibit the strongest interactions (∼5.9-fold higher than the background), the contact intensity of E-P and P-P loops is still significantly stronger (∼3-fold enrichment) than pairs between promoters and random genomic loci (**Fig. 1b**). We obtained similar results with an alternative loop calling algorithm, Chromosight^30^ (**Extended Data Fig. 1b-c**). We note that Micro-C has much higher sensitivity for detecting E-P and P-P contacts compared to Hi-C, establishing Micro-C as a more suitable assay to study genome organization relevant to transcription regulation genome-wide in an unbiased manner (**Extended Data Fig. 1d**).

**Fig 1.**
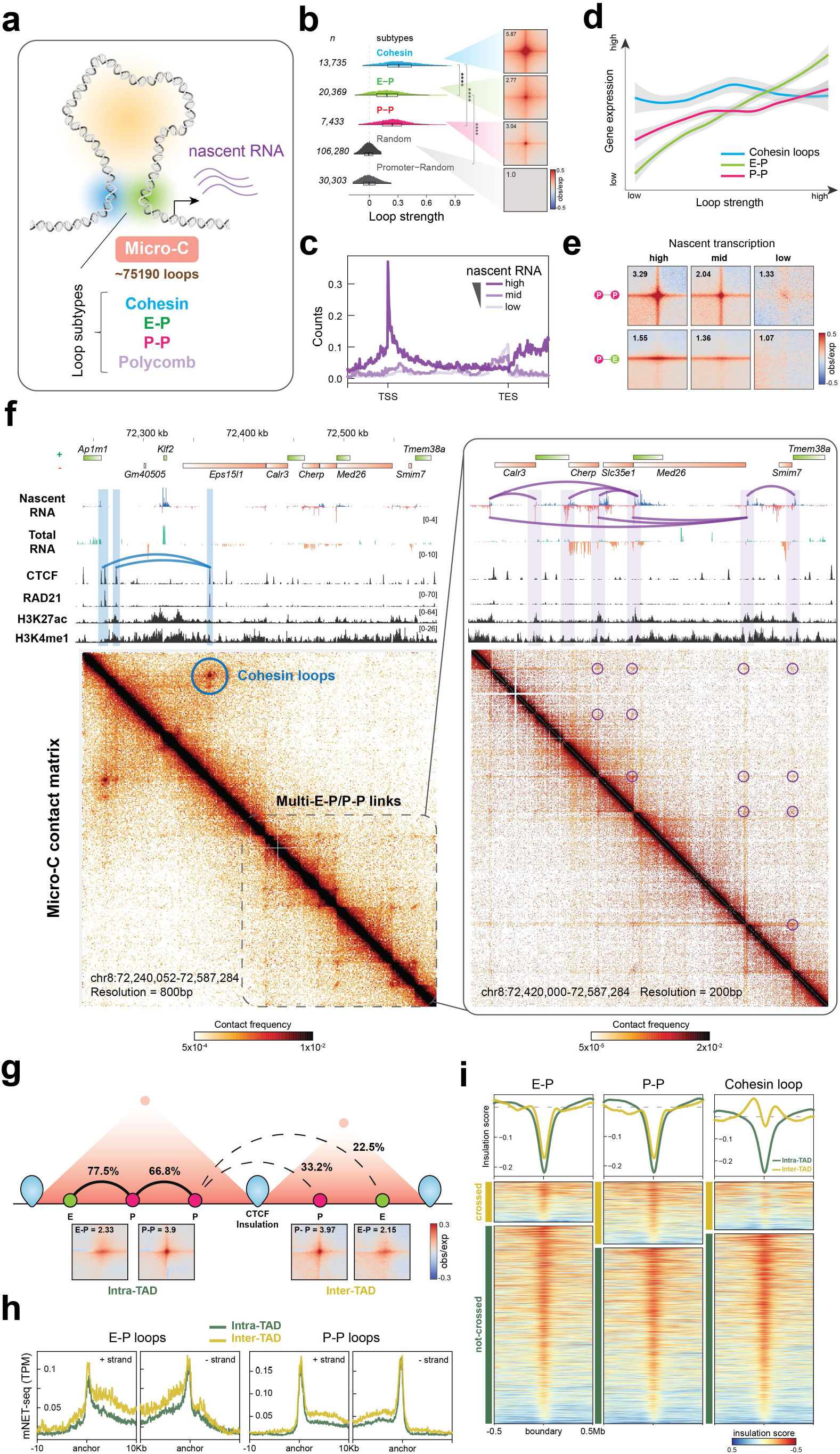
Genome-wide identification of transcription-linked chromatin loops. **a.** Micro-C identified over 75,190 chromatin loops, subclassified into four primary types (by Mustache loop caller^29^). **b.** Probability distribution of loop strength for cohesin, E-P, P-P, and random loops. The numbers of chromatin loops are shown on the left. The box plot indicates the quartiles for the loop strength score distribution. The genome-wide averaged contact signals (aggregate peak analysis (APA)) are plotted on the right. The contact map was normalized by matrix balancing and distance, with positive enrichment in red and negative signal in blue, shown as the diverging colormap with the gradient of normalized contact enrichment in log10. The ratio of contact enrichment for the center pixels is annotated within each plot. This color scheme and normalization method are used for normalized matrices throughout the manuscript unless otherwise mentioned. Asterisks denote a *p*-value < 10^−16^ by the Wilcoxon test. **c.** Genome-wide averaged transcript counts for nascent transcript profiling. Genes are grouped into high, medium, and low expression level based on nascent RNA-seq data and rescaled to the same length from TSS (transcription start site) to TES (transcription end site) on the x-axis. **d.** Rank-ordered distribution of loop strength against gene expression for cohesin, E-P, and P-P loops. The distribution for each loop type was fitted and smoothed by LOESS regression. The standard error (SE) is plotted as a gray shade along with the regression line. **e.** APA are plotted by paired E-P or P-P loops and sorted by the level of nascent transcription into high, mid, and low levels. **f.** Snapshots of Micro-C maps of a ∼300-kb region plotted with 800-bp resolution (left) and a ∼150-kb region plotted with 200-bp resolution (zoomed-in, right). The standard heatmap shows the gradient of contact intensity for a given pair of bins. This color scheme is used for Micro-C maps throughout the manuscript. Contact maps are annotated with gene boxes and 1D chromatin tracks show the signal enrichment in the same region. Features like cohesin loops (blue arched lines and circles) and E-P/P-P loops (purple arched lines and circles) enriched at stripe intersections are highlighted. **g.** Schematic (top) showing two adjacent TADs insulated by CTCF boundaries and E-P/P-P interactions either within a TAD (intra-TAD, solid arched line) or across TADs (inter-TAD, dashed arched line). TADs called by Cooltools and Arrowhead returned similar results for the ratio of boundary-crossing E-P/P-P (see Method). APA (bottom) plotted for paired E-P/P-P that either cross (inter-TAD) or do not cross (intra-TAD) a TAD boundary. **h.** Nascent transcription (± strand) at the loop anchors of intra-(green) or inter-TAD (yellow) E-P/P-P loops. **i.** Heatmap and histogram profile of insulation scores at 20-kb resolutions spanning over the 1-Mb window for intra-(green) or inter-TAD (yellow) E-P/P-P loops. Colormap shows strong insulation in red and weak insulation in blue in log10.

To gain a better understanding of the relationship between active transcription and chromatin loops, we profiled nascent transcription by mNET-seq^28^ in mESCs (**Fig. 1c and Extended Data Fig. 1e**). Newly transcribed RNAs generally have a higher correlation with E-P contacts than with compartments and TADs (**Extended Data Fig. 1f**). Specifically, the strength of E-P and P-P interactions positively correlates with the level of gene expression, while cohesin loops show no such correlation (**Fig. 1d-e**).

The region around the *Klf2* gene illustrates the complexities of fine-scale 3D genome folding and how we can use Micro-C maps to identify novel E-P contacts that are mostly unresolved in Hi-C (**Fig. 1f and Extended Data Fig. 1g**). The CTCF and cohesin ChIP-seq peaks show strong contact signals between the *Ap1m1* and *Eps15l1* genes (blue arched lines and circle), which insulate the *Klf2* gene from communicating with regions outside the loop domain. However, multiple weak interactions within the downstream 150-kb region around the *Med26* gene still occur without apparent cohesin residency at their anchors (**Fig. 1f**, right panel: purple arched lines and circles), and these contacts sharply correlate with nascent transcription signals at promoters and enhancers.

To validate these observations at a genome-wide scale, we plotted ChIP-seq data for various histone marks, transcription factors, and chromatin remodelers over the major types of loop anchors (**Extended Data Fig. 1h**). Consistent with our previous characterization of the subtypes of chromatin structures below TADs^7^ as well as a recent imaging study^31^, loop anchors enriched in CTCF and cohesin generally do not colocalize with sites of active transcription. In contrast, various transcription factors, coactivators, and Pol II are associated with E-P and P-P anchors. Thus, by coupling Micro-C with nascent RNA-seq, we can more precisely delineate which chromatin loops are associated with active transcription in a given cell type.

### E-P and P-P loops can cross TAD boundaries

TAD boundaries formed by CTCF and cohesin are thought to regulate E-P and P-P interactions in two ways: by increasing interactions inside the TAD and by blocking interactions across TADs^2^. Recent single-cell Hi-C and imaging studies have argued that TADs and their boundaries are likely to vary substantially from cell to cell^32–35^, which may allow enhancers or promoters to escape the insulation and contact regions outside of a TAD. Thus, it remains debated whether the boundaries can prevent an enhancer from interacting with and activating a gene in another TAD. Interestingly, our genome-wide analysis uncovered that, although loop interactions largely decay across distance (**Extended Data Fig. 1i**), ∼22.5% of E-P and ∼33.2% of P-P loops that cross TAD boundaries retain a comparable level of contact intensity to equidistant loops within a TAD (**Fig. 1g**). Genes located at the anchors of these inter-TAD loops also show similar or even higher expression levels in nascent or total RNA analysis (**Fig. 1h**). We postulated two possibilities that could lead to this observation: the TAD boundaries that are crossed by E-P and P-P loops have lower CTCF or cohesin occupancy or have weaker insulation propensity. We first split the TAD boundaries into two groups: “crossed” or “not crossed” by loops (**Extended Data Fig. 1j**, top). Strikingly, CTCF and RAD21 occupancy at the boundaries is almost the same regardless of whether or not the boundaries are crossed by E-P, P-P, or cohesin loops (**Extended Data Fig. 1j**, bottom). The TAD boundaries crossed by either E-P or P-P loops show only slightly weaker insulation strength than the non-crossed boundaries (**Fig. 1i**). In contrast, the boundaries that insulate the cohesin loops are substantially stronger than those that allow their crossing (**Fig. 1i**). Together, these results indicate that TAD boundaries are much more effective at insulating cohesin loops than insulating E-P or P-P loops. Over 20% of enhancers can still engage with their target sites located outside of the TAD regardless of CTCF or cohesin occupancy and insulation strength at the boundaries (**Fig. 1g**), and these boundary-crossing interactions largely correlate with transcriptional output (**Fig. 1h**). We thus hypothesize that both TAD-dependent and TAD-independent mechanisms contribute to regulation of E-P interactions.

### Acute depletion of CTCF, cohesin, and WAPL perturbs structural loops

To test whether active loop extrusion is essential for maintaining various types of chromatin loops and how these structures impact transcription, we endogenously and homozygously tagged each of the three primary loop extrusion factors (CTCF, RAD21, or WAPL) with an auxin-inducible degron (AID) by CRISPR/Cas9-mediated genome editing in mESC lines expressing the F-box protein *OsTir1* (**Fig. 2a and Extended Data Fig. 2a-b**)^36^. We confirmed near-complete degradation of the AID-tagged proteins after ∼3 hours of treatment with indole-3-acetic acid (IAA) by immunoblotting (**Fig. 2b and Extended Data Fig. 2c**). We note that previous studies employing acute CTCF or cohesin depletion used prolonged degradation (6 – 24 hours^10,11,37^), which may confound the primary molecular response with potential secondary effects^38^. To avoid such confounding effects, we chose to use 3 hours of depletion in this study.

**Fig 2.**
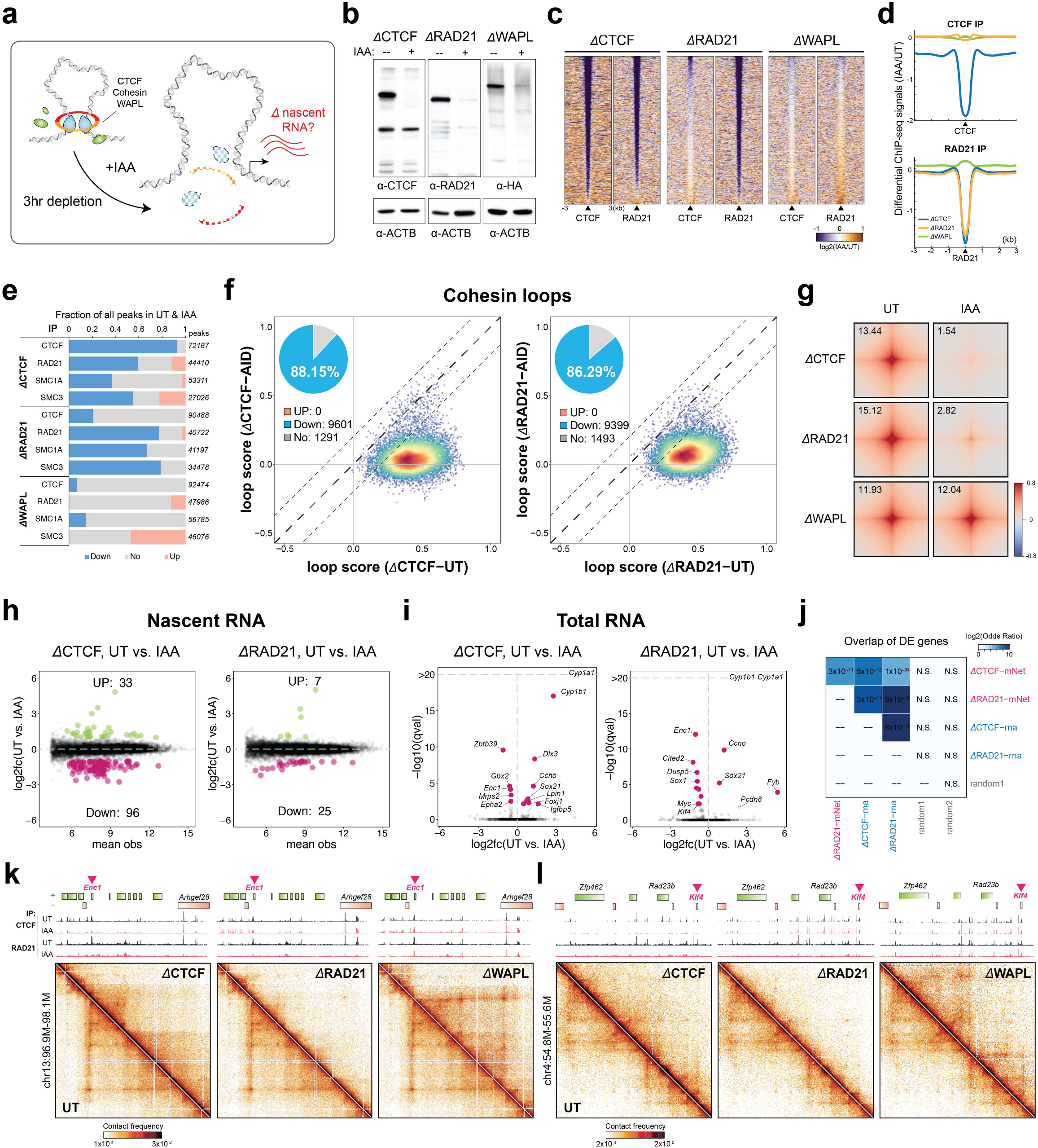
Acute depletion of loop extrusion factors affects a small set of genes. **a.** Experimental design for the degradation of CTCF, RAD21, or WAPL. **b.** Immunoblots show the degradation levels of CTCF, RAD21, WAPL, and the loading control Actin-B after 3 hours of IAA treatment. **c.** Differential ChIP-seq signals for CTCF and RAD21 in cells depleted of CTCF, RAD21, or WAPL. The peaks called by MACS2 are plotted at the center across a ±3-kb region. The colormap shows an increased signal (log2) in orange and a decreased signal in purple after IAA treatment. **d.** Histogram profile of differential ChIP-seq signals for CTCF or RAD21 in cells depleted of CTCF, RAD21, or WAPL. **e.** Summary of differential ChIP-seq peak analysis. The chart shows the fraction of down-regulated, up-regulated, or unchanged peaks after IAA treatment. The total number of peaks for each protein was summed from all peaks in untreated and IAA-treated cells. **f.** Scatter plots of loop scores for cohesin loops in the untreated and IAA-treated cells. The overlaid heatmap indicates dot density, with the highest in red and the lowest in blue. Dashed lines along the diagonal mark the range that is characterized as unchanged loops. The pie chart (inset) shows the fraction of loop intensity that is increased, decreased, or unchanged after IAA treatment. The scatter plots for comparing loop intensities between two conditions are plotted in this format throughout the manuscript unless otherwise mentioned. **g.** APA are plotted by paired cohesin peaks for the untreated and IAA-treated cells. **h.** MA plots of nascent RNA-seq for CTCF or RAD21 depletion. Differentially expressed genes (DEGs) with *q*-value < 0.05 are labeled by green (up) or pink (down). **i.** Volcano plot of total RNA-seq for CTCF or RAD21 depletion. DEGs (*q*-value < 0.01) are labeled by pink with gene name. **j.** Overlap of DEGs between different depletions and assays. Colormap shows the odds ratio in log2 and the *p*-values are annotated on the corresponding comparisons. **k-l.** Snapshots of Micro-C maps comparing chromatin interactions in the untreated (top-right) and IAA-treated (bottom-left) cells surrounding *Enc1* or *Klf4* genes. Contact maps are annotated with gene boxes and 1D chromatin tracks showing the ChIP-seq signal enrichment in the same region.

We first profiled the genome-wide binding of CTCF, RAD21, SMC1A, and SMC3 using ChIP-seq in the AID-tagged lines treated with either ethanol (untreated, UT) or IAA to degrade the tagged protein. Overall, hierarchical clustering shows high quality and reproducibility between replicates (**Extended Data Fig. 2d**). CTCF and cohesin peaks were also highly reproducible between the untreated samples (Jaccard index > 0.7) (**Extended Data Fig. 2e**). We then asked how the loss of each loop extrusion factor affects the binding of each remaining factor. Consistent with previous studies^10,39^, both CTCF and cohesin lose their occupancy after CTCF depletion (**Fig. 2c-d and Extended Data Fig. 2f-g**). Differential peak analysis^40^ confirmed that over 90% of CTCF peaks and 60% of cohesin peaks are significantly decreased upon loss of CTCF (adjusted p-value < 0.05) (**Fig. 2e and Extended Data Fig. 2h**). Thus, our results are in line with the widely accepted conclusion that CTCF is required to position cohesin^41^ and perhaps protects cohesin against release by WAPL^42^. It has been reported that cohesin is also positioned by actively transcribing Pol II and by the transcription machinery^41,43^ but the ChIP signal at these putative “CTCF-independent” loci is generally quite weak^39^(**Extended Data Fig. 2i**). Specifically, promoter-bound cohesin peaks were barely detectable in untreated cells, and though they slightly increased after CTCF depletion, we could still only detect a few thousand upregulated cohesin peaks upon CTCF degradation (**Fig. 2e and Extended Data Fig. 2i-j**)^41^. We thus suggest that in the absence of CTCF, the transcription machinery may halt cohesin extrusion but not as effectively as CTCF does. Whether the promoter-bound cohesin peaks detected after CTCF depletion are functional remains to be determined.

While cohesin peaks are generally lost upon CTCF degradation, CTCF binding is unaffected by altering the level of cohesin on chromosomes (**Fig. 2c-d**)^11^. RAD21 or WAPL depletion caused only a 10 to 20% reduction in CTCF peaks (**Fig. 2e and Extended Data Fig. 2f-g**). A recent study reported that thousands of cohesin peaks are lost near SOX2 and OCT4 binding sites after WAPL depletion^44^. Surprisingly, we did not observe even these minor changes as reported previously (**Fig. 2e and Extended Data Fig. 2h & 2k**), perhaps due to differences in degradation levels or the duration of depletion. Taken together, our ChIP-seq data confirmed effective degradation of loop extrusion factors within 3 hours and largely recapitulated previous observations^10,11,39^.

Next, we used high-resolution Micro-C to analyze the effect of CTCF, RAD21, and WAPL depletion on fine-scale 3D genome structures. Since our Micro-C data were highly reproducible across replicates (**Extended Data Fig. 3a-b**), we pooled the replicates to achieve ∼1-2 billion unique reads for each sample. At the global level, our findings largely agree with previous studies^10–12,14^. Using contact probability *P*(*s*) analysis, we observed that CTCF depletion had minimal impact on overall interactions across the genome; RAD21 depletion reduced contact frequency in the range of 10 – 200 kb but increased interactions at 300 kb – 5 Mb; and WAPL depletion showed the opposite trend, with increased contacts at 70 – 700 kb but reduced contacts at 1 – 5 Mb (**Extended Data Fig. 3c**). In addition, loop strength analysis revealed that nearly 90% of cohesin loops were lost upon depletion of CTCF or RAD21, while most loops were retained in a similar or slightly higher strength after WAPL depletion (**Fig. 2f-g and Extended Data Fig. 3d**). Indeed, after WAPL depletion, an additional ∼6,000 longer-range loops were sufficiently strengthened to meet our detection threshold (**Extended Data Fig. 3e**). These results suggest that, after CTCF depletion, cohesin-mediated DNA extrusion operates in a more unrestricted manner, resulting in the apparent loss of CTCF-anchored loops but no overall reduction of genomic interactions (**Extended Data Fig. 3c**)^10,12,41^. Interestingly, in comparison to a recent finding^45^, our data show only moderate CTCF persistence at architectural sites (e.g., loop anchors or TAD borders) upon CTCF degradation (**Extended Data Fig 3f**). CTCF binding sites associated with mitotic bookmarking^46^ and frequent occupancy sites in single-molecule foot-printing (SMF) data^47^ also did not show apparent CTCF persistence on chromatin (**Extended Data Fig 3g-h**). Given the near-complete ablation of cohesin loops after CTCF degradation (**Fig. 2g**), we suggest that the persistent residual CTCF proteins, if any, are insufficient to halt or position cohesin. Together, our Micro-C results confirmed a nearly complete disruption of cohesin loops at CTCF-anchored sites 3 hours after CTCF or RAD21 depletion and identified many emerging long-range chromatin loops in WAPL-depleted cells.

### Acute loss of CTCF, cohesin, and WAPL does not affect expression of most genes

We next asked whether acute disruption of active loop extrusion impacts the maintenance of gene expression. To capture the immediate effect of depleting loop extrusion factors on transcription, we profiled nascent transcription by mNET-seq for the control and depletion conditions along with total RNA-seq. Differential expression tests of ∼30,000 genes identified that ∼129 nascent transcripts changed in CTCF depletion, ∼32 changed in RAD21 depletion, and only 4 changed in WAPL depletion (**Fig. 2h and Extended Data Fig. 3i-j**). Total RNA-seq found ∼15 genes to be significantly deregulated in both CTCF and RAD21 depletion and ∼6 genes in WAPL depletion (**Fig. 2i and Extended Data Fig. 3j**). The differentially expressed genes (DEGs) are highly consistent between CTCF and cohesin depletion and between the assays (log2 odds ratio > 6) (**Fig. 2j**). This suggests that while CTCF and cohesin are required for the transcriptional maintenance of only a small subset of genes, those genes tend to require the presence of both factors.

Furthermore, we found that the early deregulated genes upon loss of CTCF and cohesin include many cell type-specific transcription factors (e.g., *Sox21*, *Sox1*, *Myc*, *Klf4*, *Gbx2*, *Foxj1*, *Dlx3*) (**Fig. 2i**). As expected, chromatin structures around the DEGs were strongly disrupted, often featuring loss of a boundary or domain and gain of *de novo* chromatin interactions (**Fig. 2k-l**). Since perturbing the levels of these proteins in stem cells could trigger exit from the pluripotent state and/or cell death^48–53^, we envision that a longer time of protein degradation may induce indirect, loop-independent transcriptional responses. This finding highlights the importance of distinguishing between primary and secondary effects of perturbations in the study of loop extrusion and gene expression^38^, and suggests that some conclusions from studies employing 24 or 48 hours of degradation may need to be re-evaluated^10,37,44^. In summary, we find that, while CTCF, cohesin, and WAPL may regulate some gene expression, their acute depletion affects the transcription of only a handful of genes in mESCs, which largely encode pluripotency and differentiation factors.

### Loop extrusion factors are largely dispensable for the maintenance of enhancer-promoter and promoter-promoter interactions

The very modest transcriptional changes seen upon CTCF and cohesin degradation suggest that transcriptional E-P and P-P interactions may persist for at least 3 hours after the depletion of either CTCF, cohesin, or WAPL. To test this hypothesis, we quantified the loop strength at all 75,000 dots identified in wild type mESCs in both control and loop extrusion factor depletion conditions. About 20% of loops are significantly decreased, but the remaining 60,000 loops remain largely unaltered (**Fig. 3a**). Consistent with our previous results, the disrupted loops are CTCF- or cohesin-dependent, while the persistent loops are mostly anchored by promoters and enhancers (**Fig. 3b and Extended Data Fig. 4a-b**). To further validate this, we specifically quantified the strength of loops that are anchored by E-P and P-P in the distance ranging from 5 kb to 2 Mb. Remarkably, acute depletion of CTCF and cohesin had a negligible impact on the E-P and P-P loops, with ∼80% of E-P contacts (**Fig. 3c**) and 90% of P-P contacts (**Fig. 3d**) remaining unaltered. Interestingly, although more distal loops emerge in the WAPL depleted cells (**Extended Data Fig. 3c & 3e**), the resulting higher density of cohesin on chromatin did not increase the strength of preexisting loops (**Extended Data Fig. 4c-d**). WAPL depletion also had only a minor impact on E-P and P-P interactions (**Fig. 3c-d**).

**Fig 3.**
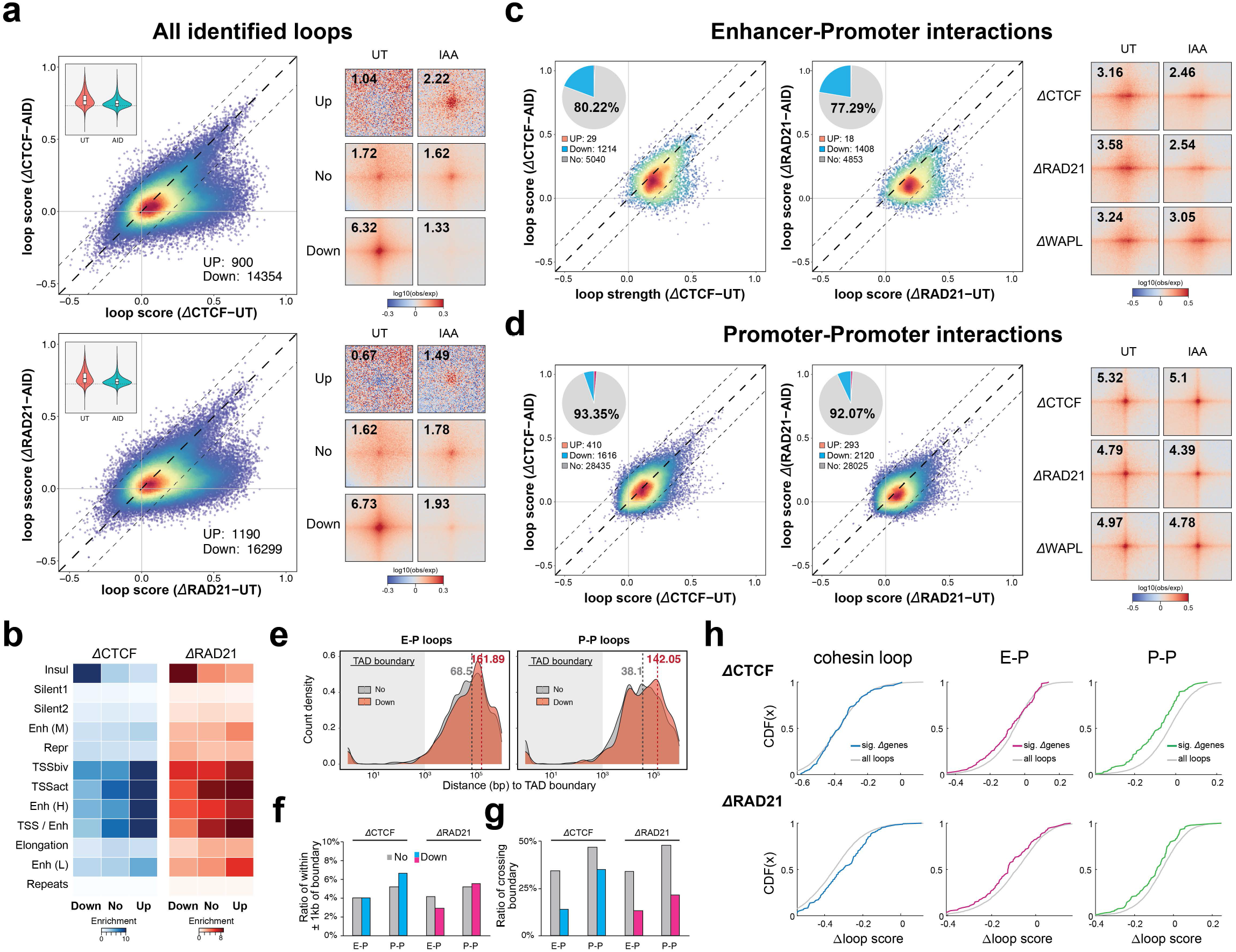
Enhancer and promoter proximity persists after the acute loss of loop extrusion factors. **a.** Scatter plots of loop scores for the called loops in the untreated and IAA-treated cells (left). The violin chart (inset) shows the distribution of loop scores for the untreated and IAA-treated conditions. APA are plotted with loops sorted by up-regulated, down-regulated, or unchanged loops (right). **b.** Enrichment of chromHMM states at loop anchors sorted by up-regulated, down-regulated, or unchanged after IAA treatment. **c-d.** Scatter plots of loop scores are plotted for paired E-P or P-P loops in the untreated and IAA-treated cells (left). APA are plotted by corresponding loop types in the untreated and IAA-treated cells (right). **e.** Length distribution of the unchanged or down-regulated E-P/P-P loops relative to TAD boundaries. **f.** Ratio of the unchanged or down-regulated E-P/P-P loop anchors located within ±1kb of TAD boundaries. **g.** Ratio of the unchanged or down-regulated E-P/P-P loop that can cross TAD boundaries. **h.** Cumulative distribution curves as a function of loop score for all loops or loop anchors overlapping with ±1kb of DEG promoters.

What are these 10-20% CTCF/cohesin-sensitive E-P or P-P loops (**Fig. 3c-d**)? According to the loop extrusion model, cohesin might either directly bridge E-P interactions at TAD boundaries, or the process of extrusion might inherently increase the frequency of long-range E-P interactions^54^. The levels of CTCF and cohesin occupancy at the anchors of these loops are much higher than those at the unaffected loop anchors (**Extended Data Fig. 4e**), suggesting that these loops may overlap with TAD boundaries, where CTCF and cohesin proteins are highly enriched. To test this possibility, we compared the genomic features of these loops relative to TAD boundaries. The lengths of the affected E-P loops are well below the average size of a TAD, ranging from 50 to 200 kb, with CTCF-sensitive E-P loops slightly larger than cohesin-sensitive loops (**Extended Data Fig. 4f**). Surprisingly, only ∼5% of the anchors are located within a 1-kb window surrounding the boundaries (**Fig. 3e-f**). Most of them are located even farther away than the unaffected loops (**Fig. 3e-f and Extended Data Fig. 4g-h**) and tend to interact with another DNA locus within the same TAD without crossing the boundary (**Fig. 3g**). We next tested whether the affected E-P or P-P interactions were associated with the DEGs in nascent RNA-seq. Indeed, E-P or P-P interactions showed a greater decrease when their associated genes were deregulated upon loss of CTCF/cohesin (**Fig. 3h**). We thus conclude that the CTCF/cohesin-sensitive E-P loops have higher CTCF and cohesin occupancy at their anchors, but these anchors are not associated with TAD boundaries. Taken together, our results suggest that E-P and P-P contacts and fine-scale gene folding largely persist and remain transcriptionally functional even after near-complete depletion of CTCF, cohesin, or WAPL.

### Probing YY1 as a candidate regulator of E-P and P-P links

Our finding that E-P and P-P interactions largely remain intact in the absence CTCF, cohesin, and WAPL suggests that other proteins are likely responsible for E-P and P-P interactions. To address this, we searched for factors specifically enriched at E-P and P-P loop anchors. BRD2, BRD4, P300, ESRRB, SP1, Mediator, YY1, pluripotency transcription factors, and chromatin remodelers are all broadly enriched at enhancer and promoter loop anchors (**Extended Data Fig. 1h**). Although Mediator has been proposed to function as a structural complex to bridge E-P interactions^55^ and promote the folding of sub-TAD structures^56^, two recent studies showed that loss of Mediator does not strongly affect E-P interactions^57,58^.

We therefore focused on YY1, a multifunctional zinc finger-containing transcription factor (**Extended Data Fig. 5a**) that is ubiquitously expressed, highly conserved, and essential for embryonic development in mammals^59^. Heterozygous YY1 mutations cause Gabriele-de Vries syndrome, which is characterized by developmental delay and intellectual disability^60^. YY1 has been implicated in chromatin looping^61^, and a previous study proposed that YY1 is a master structural regulator of E-P interactions specifically^62^. Nevertheless, upon rapid induction (1 hour) of erythroid differentiation, YY1 triggers little or no change in H3K27ac and H3K27ac-anchored HiChIP interactions^63^. These confounding results led us to investigate the role, if any, of YY1 in mediating E-P interactions using Micro-C and YY1 depletion.

### Acute YY1 depletion has little effect on global gene expression and E-P and P-P interactions

To investigate the function of YY1 in genome organization and transcriptional regulation, we fused the miniIAA7 tag^64^ to the endogenous YY1 locus to allow for rapid protein degradation. Immunoblots and flow cytometry analysis confirmed a nearly complete degradation one hour after IAA addition (**Fig. 4a and Extended Data Fig. 5a**). We chose a 3-hour degradation time for all the following assays to avoid measuring secondary effects. ChIP-seq analysis showed a clear depletion of YY1 at its cognate sites (**Fig. 4b**), which are primarily enriched at promoters, enhancers, and bivalent loop anchors (**Fig. 4c and Extended Data Fig. 5b**), consistent with its reported role in E-P interactions. The majority of YY1 peaks were diminished, but only ∼50% of peaks (n = 15075) were called significantly changed by differential peak analysis^40^ (**Fig. 4b and Extended Data Fig. 5c**). We thus only focused on those loci that had a significant loss of YY1 occupancy. We also noticed a modest decrease in cohesin occupancy after loss of YY1 (**Fig. 4b and Extended Data Fig. 5d**), which may be associated with YY1’s potential to position or halt cohesin^62^.

**Fig 4.**
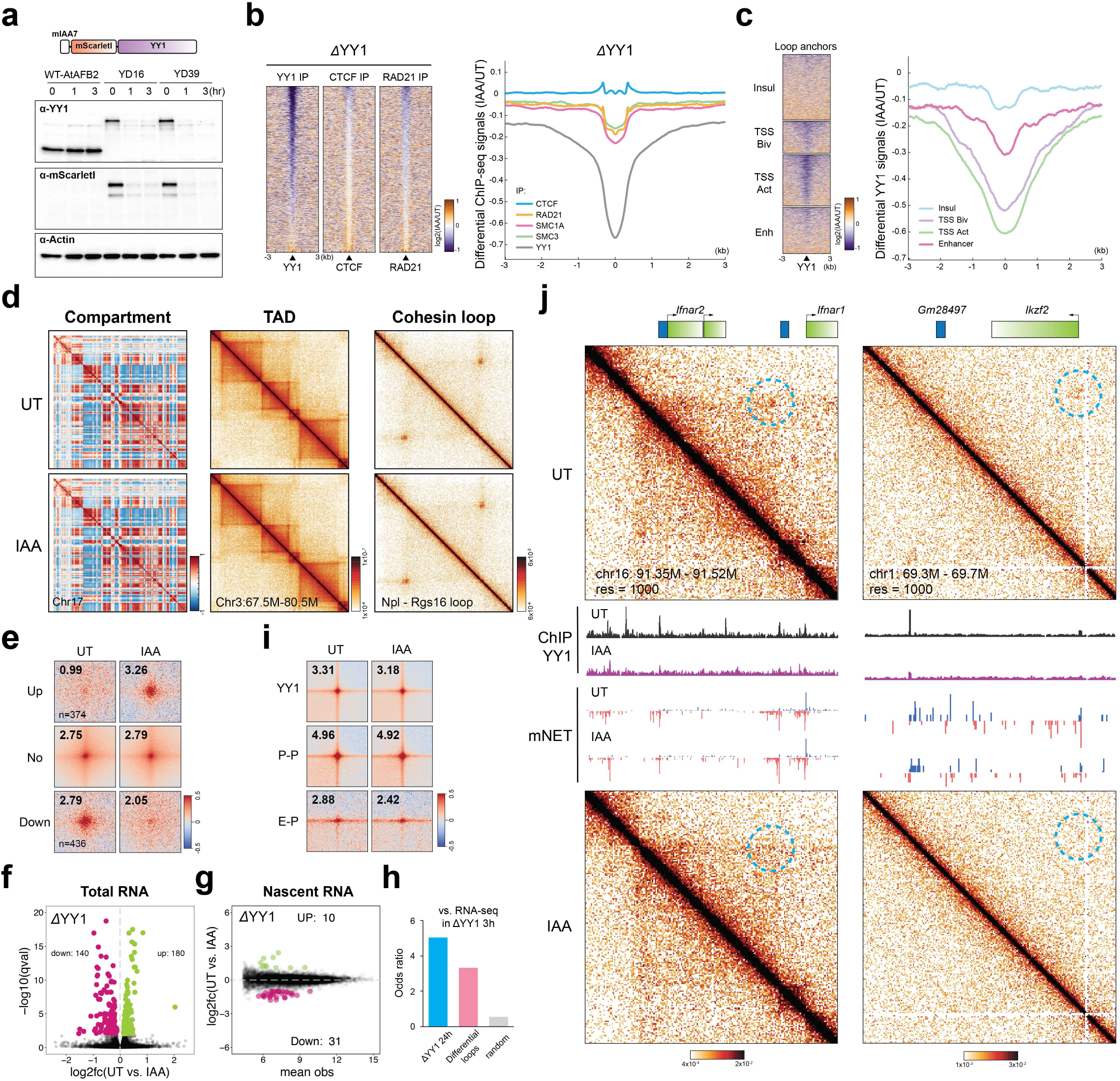
YY1 depletion does not immediately alter global gene expression and E-P or P-P proximity. **a.** Schematic for endogenous tagging for YY1 depletion and the results of immunoblots for YY1 and Actin-B. **b.** Heatmaps (left) and histogram profiles (right) of differential ChIP-seq signals for YY1, CTCF, and cohesin after YY1 depletion. **c.** Heatmaps (left) and histogram profiles (right) of differential ChIP-seq signals for YY1 around the four types of loop anchors. **d.** Overview of Micro-C contact maps at multiple resolutions in the untreated and IAA-treated cells. (left to right) Examples of Pearson’s correlation matrices showing plaid-like chromosome compartments; contact matrices showing TADs along the diagonal; contacts matrices showing cohesin loops off the diagonal. **e.** APA are plotted with loops sorted by up-regulated, down-regulated, or unchanged. **f.** Volcano plot of total RNA-seq for YY1 depletion. DEGs (*q*-value < 0.01) are colored by green (up) or pink (down). **g.** MA plots of nascent RNA-seq for YY1 depletion. DEGs with *q*-value < 0.05 are colored by green (up) or pink (down). **h.** Overlap of DEGs in RNA-seq after YY1 depletion with 1) RNA-seq after 24-hour YY1 depletion, 2) genes associated with differential loops, and 3) a set of random genes. **i.** APA are plotted by corresponding loop types in the untreated and IAA-treated cells. **j.** Snapshots of Micro-C maps comparing chromatin interactions in the untreated (top) and IAA-treated (bottom) cells surrounding the *Ifnar2* or *Ikzf2* genes. Contact maps are annotated with gene boxes and genome browser tracks showing YY1 ChIP-seq signal enrichment and mNET-seq signals with the plus strand in blue and the negative strand in red.

To characterize YY1’s role in 3D genome organization, we acquired ∼850 M unique Micro-C reads after pooling high-quality replicates from mock-treated and YY1-depleted cells (**Extended Data Fig. 5e**). We found that YY1 depletion has no strong effect on chromatin compartments, TADs, and cohesin loops (**Fig. 4d and Extended Data Fig. 5f**), and no significant change in genome-wide chromatin interaction probabilities (**Extended Data Fig. 5g**). YY1 was proposed to be a causally required structural regulator of transcription and E-P interactions^62^. Surprisingly, acute removal of YY1 only significantly affected ∼800 loops (**Fig. 4e**), and ∼325 and ∼40 genes in the RNA-seq and mNET-seq profiling, respectively (**Fig. 4f-g and Extended Data Fig. 5h**). The DEGs in RNA-seq data significantly overlapped with the genes associated with differential loops or DEGs identified after 24 hrs of YY1 depletion^62^ (**Fig. 4h and Extended Data Fig. 5h**), indicating a high correlation between E-P loops and gene expression, as well as confirming capture of an early response to YY1 degradation. More importantly, genome-wide pileup analysis for the intersections of YY1 peaks, E-P loops, and P-P loops showed only a very minor change in loop intensity after YY1 depletion (**Fig. 4i and Extended Data Fig. 5f**). This result suggests that the maintenance of most E-P loops and their regulatory functions in general do not require the presence of YY1, at least within a 3-hour depletion window. Nevertheless, a specific set of loci appears to require the presence of YY1 in order to interact with their cis-regulatory elements. For example, at the *Ifnar2* and *Ikzf2* gene loci, the weak E-P or P-P interactions were lost upon YY1 depletion and were often accompanied with either an increase or decrease in nascent transcription (**Fig. 4j and Extended Data Fig. 5i**). Taken together, although YY1 may be required for some limited set of E-P and P-P interactions, these results are not consistent with the model that YY1 is a general master structural regulator of E-P interactions in mESCs as previously proposed^62^.

### Single-molecule imaging reveals YY1 binding dynamics and nuclear organization

The surprisingly meager effects of YY1 on chromatin looping, E-P, and P-P interactions might result from YY1 DNA binding being very transient and/or due to only a small fraction of YY1 proteins being bound to DNA in live cells. To better understand the dynamics and mechanisms underlying YY1 function in living cells, we tagged YY1 with HaloTag, a self-labeling protein tag that can be covalently conjugated with cell-permeable synthetic dyes suitable for single-molecule imaging in live cells (**Extended Data Fig. 6a**)^65^. We either knocked in a HaloTag at the N-terminus of the endogenous YY1 by CRISPR-Cas9-mediated genome editing (**Fig. 5a and Extended Data Fig. 6b**) or ectopically expressed YY1 fused with HaloTag via a minimal mammalian *L30* promoter by *PiggyBac* transposition. We isolated and validated two homozygously tagged clones, designated YN11 and YN31. We further confirmed that the Halo-tagged YY1 protein in both endogenously tagged and ectopically expressing clones was of the correct size and expressed at close to endogenous levels (**Fig. 5a and Extended Data Fig. 6c**). Live-cell confocal imaging validated that HaloTag-YY1 was predominantly localized within the nucleus and appeared to be non-homogeneously distributed throughout the nucleoplasm, with noticeable exclusion from nucleoli (**Fig. 5b-c and Extended Data Fig. 6d**). We observed bright YY1 puncta sporadically clustered within nucleoli, which is reminiscent of the nucleolar localization of TBP and its function on Pol III transcription (**Fig. 5b-c and Extended Data Fig. 6d**)^66,67^. We then visualized the nuclear distribution of YY1 at single-molecule resolution by using photoactivated localization microscopy (PALM) (**Fig. 5d**). YY1 displays a punctate distribution with many high-density clusters throughout the nucleoplasm. These high-density clusters are unlikely to be a fixation artifact, since such puncta were also visible in Airyscan live-cell imaging (**Fig. 5c and Extended Data Fig. 6d**). YY1 has been thought to be evicted from chromosomes during mitosis in fixed-cell imaging experiments^68^. However, our live-cell imaging showed continued YY1 residency on mitotic chromosomes, suggesting that YY1 may be involved in mitotic bookmarking (**Fig. 5b and Extended Data Fig. 6d**)^69^. Together, these results validate our homozygous HaloTag-YY1 knock-in cell lines and reveal that YY1 binds mitotic chromosomes and forms local high concentration hubs in the nucleus.

**Fig 5.**
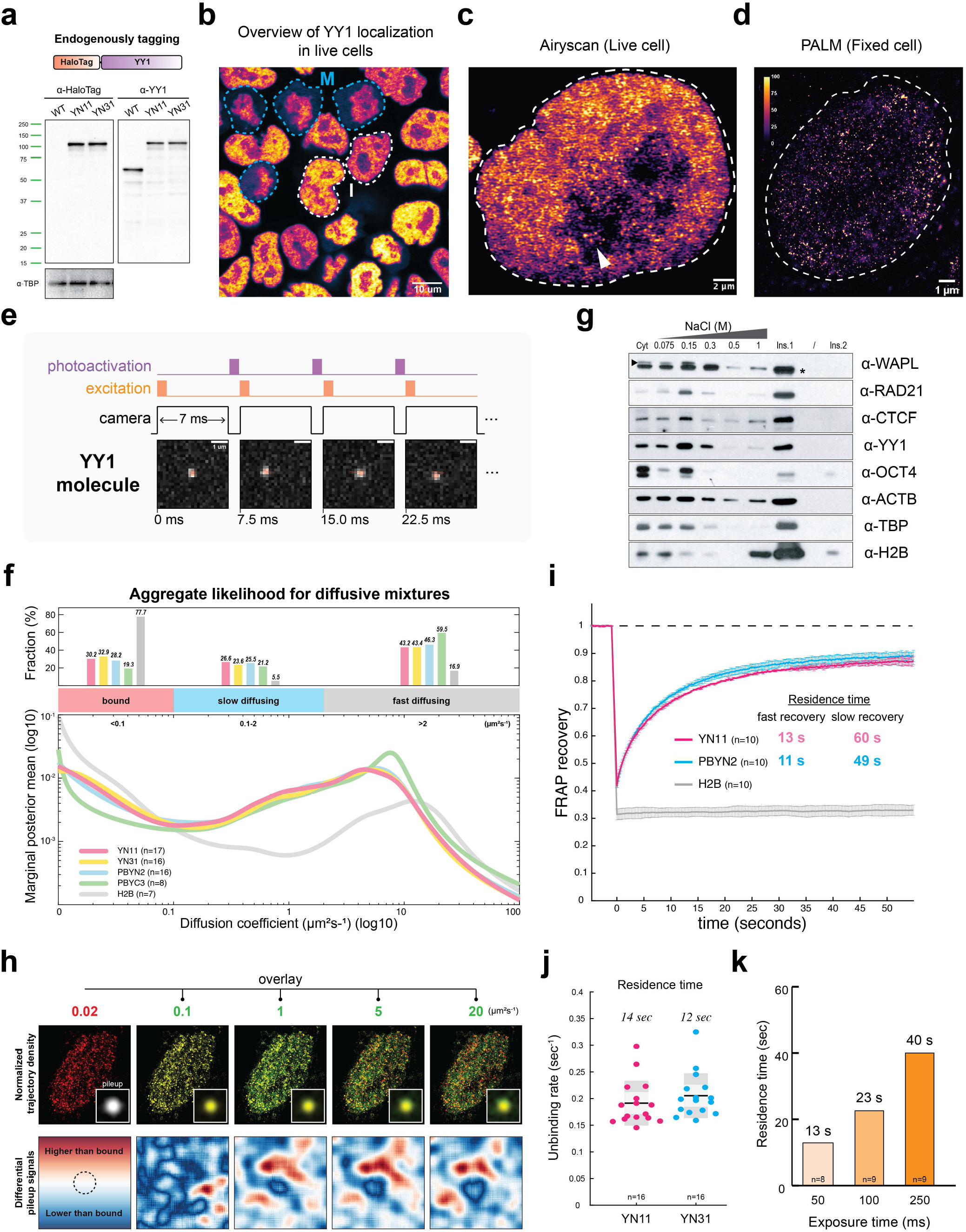
Dynamics of YY1 protein binding. **a.** Schematic for endogenously tagging YY1 with HaloTag and the results of immunoblots for YY1 and TBP. **b.** Live-cell confocal imaging for HaloTag-YY1 stained with 500 nM TMR. White dashed lines label cells in interphase and blue dashed lines label mitotic cells. Scale bar: 10 µm. **c.** Airyscan-resolved live-cell confocal imaging for YY1 (n=13). Arrow points to sporadic loci within the nucleolus. Scale bar: 2 µm. **d.** PALM imaging for YY1 (n=30). Colormaps color the signal ranging from 0-100. Scale bar: 1 µm. **e.** Overview of spaSPT with illumination patter and representative raw images for YY1 with tracking overlaid. Scale bar=1 µm. **f.** The aggregate likelihood for diffusive YY1 molecules. The bar graph (top) shows the fractions of YY1 binned into bound, slow, and fast diffusing subpopulations. (bottom) Estimation of YY1 diffusion coefficients by regular Brownian motion with marginalized localization errors. **g.** Immunoblots of proteins after a series of salt extractions. Cyt=cytoplasmic fraction; Ins=Insoluble fraction. **h.** Spatial reconstruction of spaSPT data for YY1’s trajectory densities. YY1 trajectories are binned by diffusion coefficients as indicated. The bound trajectories are colored in red. The diffusing trajectories are colored in green and overlaid with the bound trajectories. The insets show the averaged signals for all identified clusters aggregating at the center of the plot with the same imaging overlays. Differential signals comparing each diffusing fraction to the bound fraction are plotted at the bottom. The divergent colormap shows a higher signal than the bound in red and lower than the bound in blue. **i.** FRAP analysis of YY1 bleached with a square spot. Error bars indicate the standard deviation of each acquired data point. **j.** Slow-SPT for measuring YY1’s residence time (exposure time = 100 ms). Each data point indicates the unbinding rate of YY1 molecules in a single cell. The box plot shows the quartiles of data. **k.** Slow-SPT for measuring YY1’s residence time with multiple exposure times.

Having characterized our cell lines, we next interrogated YY1 protein dynamics and target search mechanisms. We took advantage of the stroboscopic photo-activation single-particle tracking technique (spaSPT)^70,71^ to minimize motion-blur and tracking errors to unambiguously trace the movement of individual YY1 molecules at a frame rate of ∼133 Hz (**Fig. 5e and Extended Data Fig. 6e**). Spots corresponding to HaloTag-YY1 molecules were detected and tracked to form trajectories^72^. We then inferred distributions of diffusion coefficients from the spaSPT data using a finite-state approximation to the Dirichlet process mixture mode (State Array) implemented in the newly developed package Spagl (**Extended Data Fig. 6e-f**)^72^. The diffusive mixtures can be attributed to at least two major apparent diffusion states^72^, including a bound population (diffusion coefficient (*D_bound_*) < 0.1 µm²s^-^¹) and a broad mixture of freely diffusing molecules (*D_free_* > 0.1 µm²s^-^¹) (**Fig. 5f**). We found that ∼31% of YY1 is in an immobile state, presumably bound to chromatin, with the remaining population exhibiting either slow diffusion (∼26%; *D_slow_* ∼ 0.1 – 2 µm²s^-^¹) or fast diffusion in the nucleoplasm (∼43%; *D_fast_* > 2 µm²s^-^¹) (**Fig. 5f**). These measurements largely agree with kinetic modeling of displacements obtained with the Spot-On algorithm (**Extended Data Fig. 6g**)^71^. As an independent and orthogonal validation of these results, we biochemically probed YY1 chromatin binding affinity by subjecting freshly isolated nuclei to a series of washes with increasing salt concentrations (**Extended Data Fig. 7a**). The majority of YY1 is extracted at low salt concentrations (75 – 300 mM), but a subpopulation stayed bound on chromatin (∼28%), resisting 1 M washes, consistent with the ∼31% chromatin bound population estimated by spaSPT. The fraction of YY1 stably associating with chromatin is substantially lower than CTCF (∼43%) and cohesin (∼65%) (**Fig. 5g and Extended Data Fig. 7b**).

To further link the spatial relationship of YY1 proteins with their different diffusion rates, we reconstructed the spatial distributions of trajectories by their likelihood of diffusion coefficients at 0.02, 0.1, 1, 5, and 20 µm²s^-^¹. Consistent with the live-cell imaging and PALM results, the immobile fraction showed apparent cluster-like structures with ∼12 – 30 trajectories per spot (from a total of ∼15,000 trajectories per cell), while faster moving YY1 subpopulations showed weaker propensity to cluster within a constrained area (**Extended Data Fig. 7c-d**). Overlays of the immobile trajectories with the other fractions showed that the molecules within the bound regime had nearly complete overlap, but the faster-diffusing molecules were less likely to co-localize with the immobile clusters (**Fig. 5h**). The averaged (**Fig. 5h, insets**) and differential cluster signals (**Fig. 5h, bottom panel**) between the immobile and the other fractions (**Extended Data Fig. 7c-d**) further confirmed that the faster-moving molecules more frequently travel to areas in the vicinity of the immobile clusters. These results are consistent with chromatin-bound YY1 proteins forming clusters, while the diffusing YY1 molecules appear to search the 3D nuclear space outside of the clusters. We previously made similar observations for CTCF^27,70,73^.

The residence times of transcription factors bound at their targets often correlate with their functional outcomes^74–76^. To estimate the overall residence time of the bound fraction of YY1, we used fluorescence recovery after photobleaching (FRAP) to measure *in vivo* protein binding kinetics by fitting the fluorescence recovery curve to a kinetic model^77,78^. Using a reaction-dominant FRAP model, we estimated a residence time of ∼13 seconds for the majority of YY1 molecules (**Fig. 5i and Extended Data Fig. 7e-f**). Interestingly, we noticed that ∼5-10% of YY1 bound chromatin for a longer period of time with an average residence time of ∼50 seconds (**Fig. 5i and Extended Data Fig. 7e-f**). We also employed slow-SPT as an orthogonal approach to measure YY1 residence times. For slow-SPT, we imaged individual molecules at an exposure time of 100 ms, which blurs fast-moving molecules into the background and effectively captures stable binding events^79^. Model fitting for the survival curve of molecules^79^ yielded a residence time of ∼13 seconds for YY1 (**Fig. 5j**). Consistent with the FRAP data showing ∼5-10% of long binders, slow-SPT with exposure times from 50 to 250 ms^80^ further revealed a subpopulation of YY1 that binds chromatin for over half a minute (**Fig. 5h**).

While YY1’s average residence time of ∼13-40 seconds is similar to that of many transcription factors, it is much shorter than the residence times of known structural factors such as CTCF (∼1-4 min) and cohesin in G1 (∼20-25 min) in mESCs^70^. These results may explain why YY1 depletion has a marginal effect on chromatin looping than does CTCF and cohesin depletion. CTCF/cohesin loops are generally stronger and almost completely lost upon CTCF/cohesin depletion (**Fig. 2f-g**), whereas YY1 loops tend to be weaker and less affected by YY1 depletion (**Fig. 4e&i**). Furthermore, the chromatin bound fraction of YY1 (∼30%) is considerably smaller than that for CTCF (∼50-60%)^70^. Taken together, our imaging experiments suggest that a smaller fraction of YY1 is bound to DNA and that YY1 binding is more dynamic than CTCF, which may help explain why YY1 protein depletion has a much weaker effect on looping and 3D genome folding.

### Cohesin depletion alters YY1’s chromatin binding

We recently showed that CTCF clusters are enriched in diffusive CTCF proteins near their binding sites, thereby accelerating their target search^73^. To test if CTCF and cohesin may similarly affect YY1’s target search, we endogenously fused an auxin-inducible degron (AID) to CTCF or RAD21 in the HaloTag-YY1 parental line and confirmed > 90% depletion after 3 hours of IAA treatment (**Fig. 6a and Extended Data Fig. 8a**). Despite the high degradation efficiency, neither YY1’s nuclear distribution nor its clustering was strongly affected after acute loss of CTCF and cohesin in either live- or fixed-cells (**Fig. 6b-c and Extended Data Fig. 8b**). This suggests that the maintenance of YY1 hubs is independent of CTCF and cohesin.

**Fig 6.**
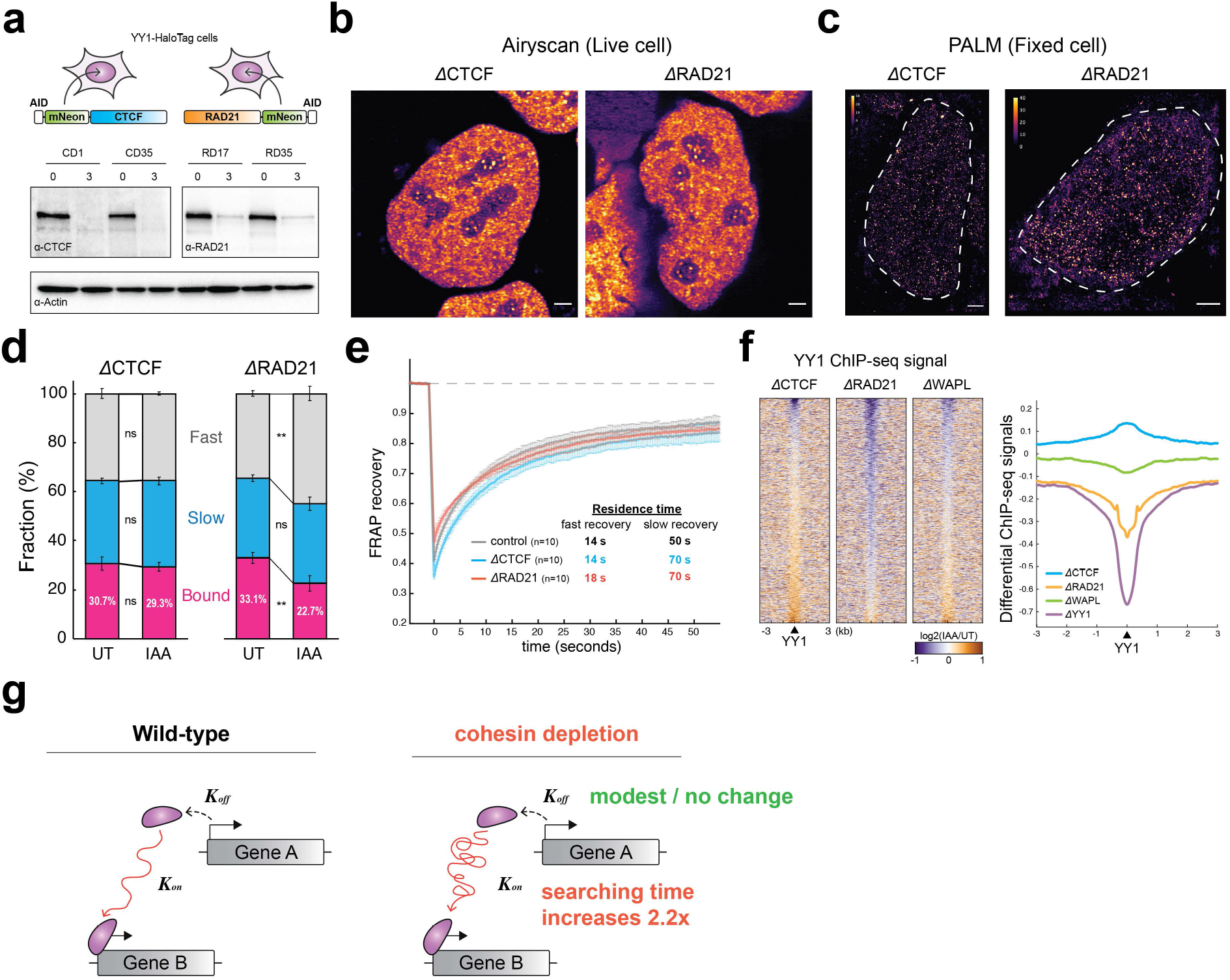
Cohesin depletion alters YY1’s target search efficiency. **a.** Schematics for endogenously tagging CTCF or cohesin with AID in the HaloTag-YY1 cell line (clone YN11) and immunoblots of CTCF, RAD21, and Actin-B. **b.** Airyscan-resolved live-cell confocal imaging for HaloTag-YY1 stained with 500 nM TMR in CTCF- or RAD21-depleted cells (n=6 for each depletion). Scale bar: 1 µm. **c.** PALM imaging for YY1 (n=13 for each depletion). Colormaps color the signal ranging from 0-40. Scale bar: 1 µm. **d.** The stacked bar graph shows the fractions of bound, slow, and fast diffusing YY1 in the untreated and IAA-treated cells, obtained by Spagl analysis. **e.** FRAP analysis of YY1 in the control, CTCF-depleted, or RAD21-depleted cells. **f.** Heatmaps (left) and histogram profiles (right) of differential ChIP-seq signals for YY1 after CTCF, RAD21, or WAPL depletion. **g.** Dynamic model of how cohesin or cohesin-mediated structures may regulate TF target search.

We next examined YY1 nuclear target search efficiency in the absence of CTCF and cohesin using spaSPT. Although CTCF depletion had no major effect, cohesin depletion resulted in a modest but reproducible decrease from ∼33% to 22% (∼31% decrease; *p*-value < 0.01) in the bound fraction of YY1 (**Fig. 6d and Extended Data Fig. 8c**). A lower bound fraction could either result from a shorter residence time (*k*_off_) or slower target search (*k*_on_). To distinguish between these possibilities, we analyzed the FRAP data and found YY1 residence times (*k*_off_) to be only weakly affected by CTCF and cohesin depletion (**Fig. 6e**). We therefore conclude that cohesin loss may affect YY1 target search (*k*_on_). Specifically, we estimated a ∼54% decrease in *k_on_* after cohesin depletion, resulting in a ∼2.2-fold longer YY1 search time (UT=28 s; IAA=61 s), the time it takes YY1 on average to find and bind a cognate binding site after dissociating from DNA.

To independently test this SPT finding, we analyzed our ChIP-seq data. We found that ∼3505 YY1 peaks (total peaks = ∼41989) were lost after RAD21 degradation, and over 82% of these loci were associated with promoter regions (**Fig. 6f and Extended Data Fig. 8d-e**). In contrast, both CTCF and WAPL depletion had a negligible effect on YY1 occupancy (**Fig. 6f and Extended Data Fig. 8d-e**). These differences suggest that cohesin likely facilitates YY1 chromatin binding, especially at promoter regions.

Taken together, our results reveal a previously unappreciated role for cohesin in accelerating the target search of transcription factors resulting in increased YY1 chromatin binding as measured by SPT, FRAP, and ChIP-seq. These findings suggest that long-term cohesin depletion experiments must be interpreted with caution since cohesin depletion results in both direct and indirect effects including diminished general transcription factor binding to DNA (**Fig. 6g**).

## Discussion

Both the extent and mechanism by which CTCF and cohesin-mediated loop extrusion regulates transcription has remained puzzling and hotly debated^10,11,22,23,31,34,37,41,45,81^. Previous work has shown that CTCF or cohesin depletion abolishes TADs genome-wide and modestly affects gene expression^10–14^. However, since Hi-C cannot readily capture fine-scale transcriptionally relevant E-P and P-P interactions, how E-P and P-P interactions are regulated has remained unclear. Here we applied high-resolution Micro-C to overcome this limitation. Surprisingly, we found that neither CTCF, cohesin, WAPL, nor YY1 are required for the maintenance of E-P or P-P loops or transcription at least within a 3-hour window after depletion. Our findings, together with other evidence, allow us to distinguish and/or eliminate several models of E-P interactions previously assigned to these ubiquitous structural proteins (**Fig. 7**).

**Fig 7.**
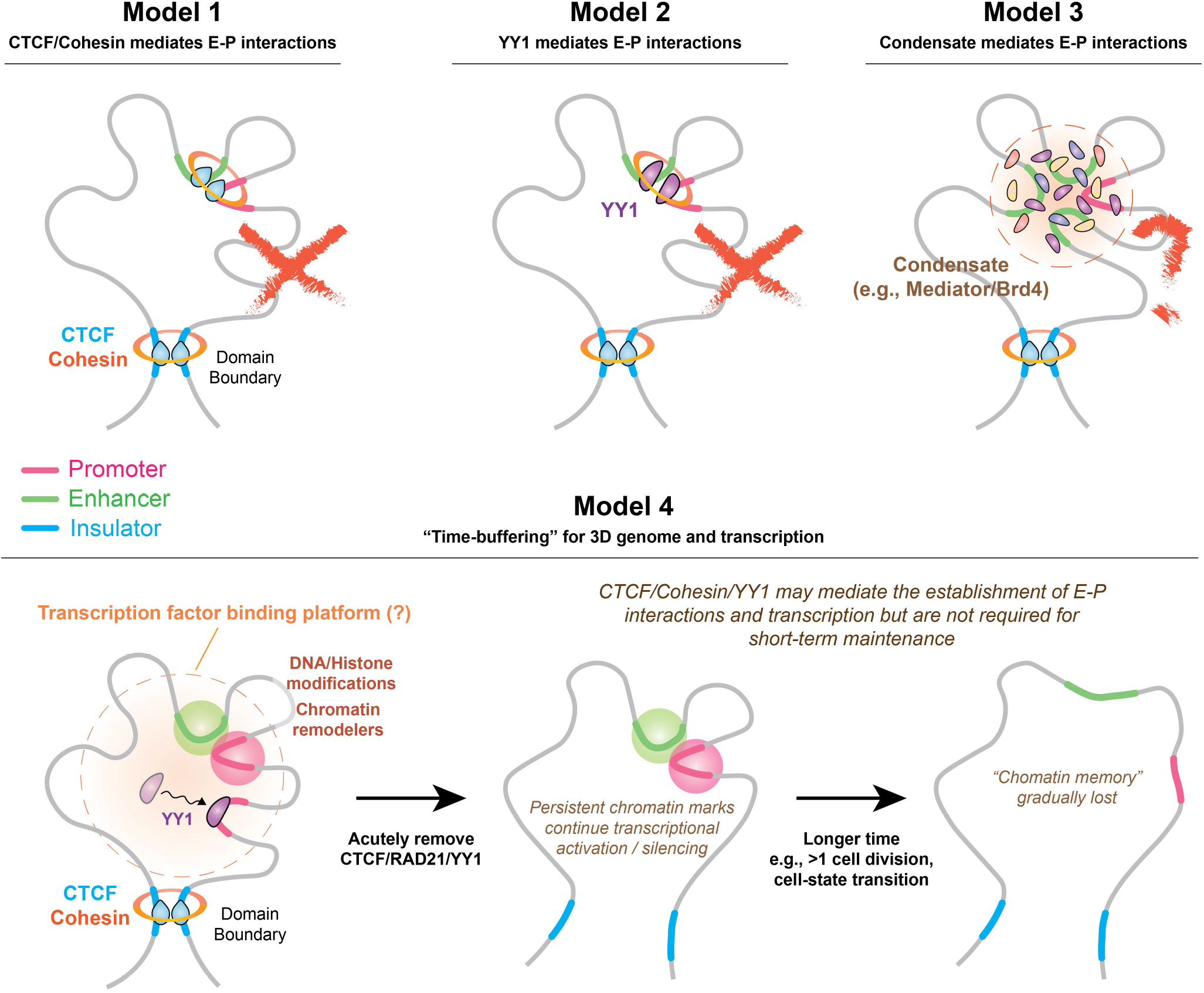
Models of enhancer-promoter interactions and transcription in the context of 3D genome organization.

First, CTCF and cohesin have been proposed to either directly bridge E-P interactions^55^ or to indirectly mediate E-P interactions by increasing contact frequency inside TADs (**Fig. 7, Model 1**)^82^. Our finding that acute CTCF, cohesin, and WAPL depletion minimally affects gene expression (**Fig. 2h-j**) and E-P interactions (**Fig. 3**) disfavors this model for short-term maintenance of E-P interactions, though CTCF and cohesin may still help establish E-P interactions indirectly. We propose that loop extrusion may often be a separable mechanism from most E-P interactions and transcription, which is further supported by the following observations: 1) Over 20% of E-P or P-P loops can cross TAD boundaries and retain high contact probability and transcriptional activity (**Fig. 1**); 2) Only a very small handful of genes showed altered expression levels after CTCF, cohesin, or WAPL depletion (**Fig. 2**)^10–14^; 3) The majority of E-P or P-P loops persists after depletion of these structural proteins (**Fig. 3**)^37,81^; 4) CTCF/cohesin generally do not colocalize with transcription loci^31^; 5) E-P loops and transcription can be established in some cases even with no CTCF/cohesin expression^22,23,83^. Second, YY1 was proposed to be a master structural regulator of E-P interactions^62^ (**Fig. 7, Model 2**). However, our Micro-C data is inconsistent with this model for the maintenance of E-P interactions, since acute YY1 depletion has little effect on E-P and P-P interactions or gene expression. Third, phase-separated transcriptional condensates have been proposed to mediate gene activation and E-P interactions^84^. Pol II, Mediator, BRD4, and many transcription factors containing intrinsically disordered regions (IDRs) that tend to aggregate into local-high concentration hubs in the nucleus^85–87^. Hub formation is thought to be a general property of TFs used to engage with regulatory elements and keep them in the spatial vicinity for gene regulation^84–87^. While not specifically addressed in this study, recent studies employing acute depletion/inhibition of Mediator and BRD4, which are supposed to be key players in condensate formation, also found no drastic effects on E-P interactions^57,58,88^. Thus, condensates also do not seem to be generally required for the maintenance of E-P interactions (**Fig. 7, Model 3**). In summary, we conclude that neither transcriptional condensates, CTCF, cohesin, WAPL, nor YY1 are generally required for the short-term maintenance of E-P interactions and the subsequent expression of most genes after acute depletion and loss of function.

Does this mean that none of these factors play any role in regulating E-P interactions and gene expression? The evidence that CTCF and cohesin can directly or indirectly regulate E-P interactions and affect gene expression in many cases is overwhelming (see examples below). To reconcile these studies with our new observations, we propose a “time-buffering” model (**Fig. 7, Model 4**). In this model, CTCF, cohesin, and architectural factors contribute to the establishment of E-P interactions, but not to their maintenance. Instead, once established, a molecular memory (e.g., histone modifications^89^, chromatin remodeling^90–92^, DNA modification^93–95^, lncRNAs^96,97^, etc.) may be sufficient to maintain E-P interactions and gene expression for several hours without the contribution of CTCF, cohesin, and other architectural factors. We propose that this time-buffering model reconciles our observations with the unambiguous genetic evidence that CTCF and cohesin regulate some E-P interactions. This evidence includes the following: 1) Insertion of CTCF sites between an enhancer and a promoter can both reduce E-P interactions and strongly reduce gene expression^98–100^; 2) CTCF binding site silencing^101,102^ or genetic CTCF binding site loss^21,103,104^ can cause aberrant E-P interactions and gene expression and drive disease; 3) Inversion or repositioning of CTCF sites can redirect E-P interactions that cause gene misexpression and diseases^105,106^. Two recent studies have also proposed variants of a time-buffering model based on mathematical modeling of E-P interactions and gene expression^100,107^. In both models, individual E-P interactions are memorized – either as long-lived promoter states^100^ or as long-lived “promoter tags”^107^ – such that gene expression can be temporally uncoupled from E-P interactions, yet still be causally linked. An alternative, more conservative interpretation of our data and the evidence cited above is that CTCF and cohesin only regulate a very small, unique sets of genes in specific biological processes, and their effect on a handful of loci simply cannot be generalized as a universal rule. In summary, we propose a time-buffering model for E-P interactions where CTCF, cohesin, and architectural factors contribute to establishing E-P interactions at some genes, but where these factors are not generally required for the maintenance of E-P interactions and gene expression, except for a very small number of pluripotency genes in mESCs.

Here we also provide the first comprehensive study of YY1 dynamics and nuclear organization (**Fig. 5**). We discovered that: 1) YY1 binds mitotic chromosomes, implicating its potential function for mitotic bookmarking or transcriptional memory through the cell cycle; 2) YY1 forms nuclear hubs similar to CTCF and cohesin^27,70,73^ that may associate with E-P or P-P interactions; 3) About 31% of YY1 proteins are bound to chromatin with an average residence time of ∼13-40 seconds. Consistent with lower and less stable YY1 chromatin binding compared to CTCF and cohesin (CTCF: ∼50% bound for ∼1-4 min; Cohesin ∼40-50% bound for ∼20-25 min)^70^, acute YY1 depletion has little, if any, effect on 3D genome folding compared to CTCF and cohesin depletion.

Surprisingly, we found that cohesin depletion, but not CTCF depletion, significantly reduces YY1 chromatin binding and increases its target search time from 28 to 61 seconds. While more work on other transcription factors (TF) will be necessary to test the generality of this model, we propose that cohesin or its associated chromatin structures could serve as a general scaffold or platform for TFs that facilitates and increases TF binding to chromatin. Interestingly, the subunits of cohesin as well as its loading and unloading complexes are composed of multiple segments of IDRs, which have the potential to establish weak protein-protein interactions or multivalent interactions. These weak transient interactions may further engage with IDRs in TFs and thus facilitate their binding to chromatin^108,109^. Although more quantitative works will be necessary to unveil these mechanisms, in addition to its roles in loop extrusion, DNA repair, replication, and chromosome segregation, cohesin might also serve as a general scaffold that facilitates TF binding to chromatin and could be critical for ensuring the precise timing of gene activation and silencing during embryonic development and cell-state transitions^83^.

In summary, we have comprehensively investigated the role of CTCF, RAD21, WAPL, and YY1 in finer-scale chromatin structure, nascent transcription, as well as YY1 dynamics and nuclear organization. We propose a time-buffering model, where architectural proteins generally contribute to the establishment, but not the short-term maintenance, of E-P interactions and gene expression, and we also propose that cohesin plays an underappreciated role as a general scaffold that could facilitate TF binding to chromatin. The connection linking protein dynamics to chromatin structure opens a new avenue to rethink the mechanism of transcriptional regulation in the context of 3D genome organization.

## Acknowledgment

T.S.H., C.C., and E.S. conceived and designed the project. C.C. performed all biochemical and ChIP-seq experiments and analysis with E.S.’s assistance. E.S. generated all plasmids and cell lines with T.S.H.’s and Gina M. Dailey’s assistance. C.C. and A.S.H. generated the parental degradation cell lines. T.S.H. performed Micro-C assays and analysis. E.S. and T.S.H. performed all imaging experiments and analyses. T.S.H. drafted the manuscript, and C.C., E.S., A.S.H., R.T. edited the manuscript. R.T. and X.D. supervised the project. Funding: This work was supported by the California Institute of Regenerative Medicine grant LA1-08013 (X.D.), the Koret UC Berkeley-Tel Aviv University Initiative grant (T.S.H. & X.D.), and by the Howard Hughes Medical Institute (003061, R.T.). T.S.H. is a postdoc fellow of the Koret UC Berkeley-Tel Aviv University Initiative. E.S. was an undergraduate fellow of SURF Rose Hills Independent at UC Berkeley. A.S.H. acknowledges support from the National Institutes of Health under Grant R00GM130896. We thank all members of the Tjian and Darzacq lab for comments on the manuscript.

---

**Extended Data Fig 1.**
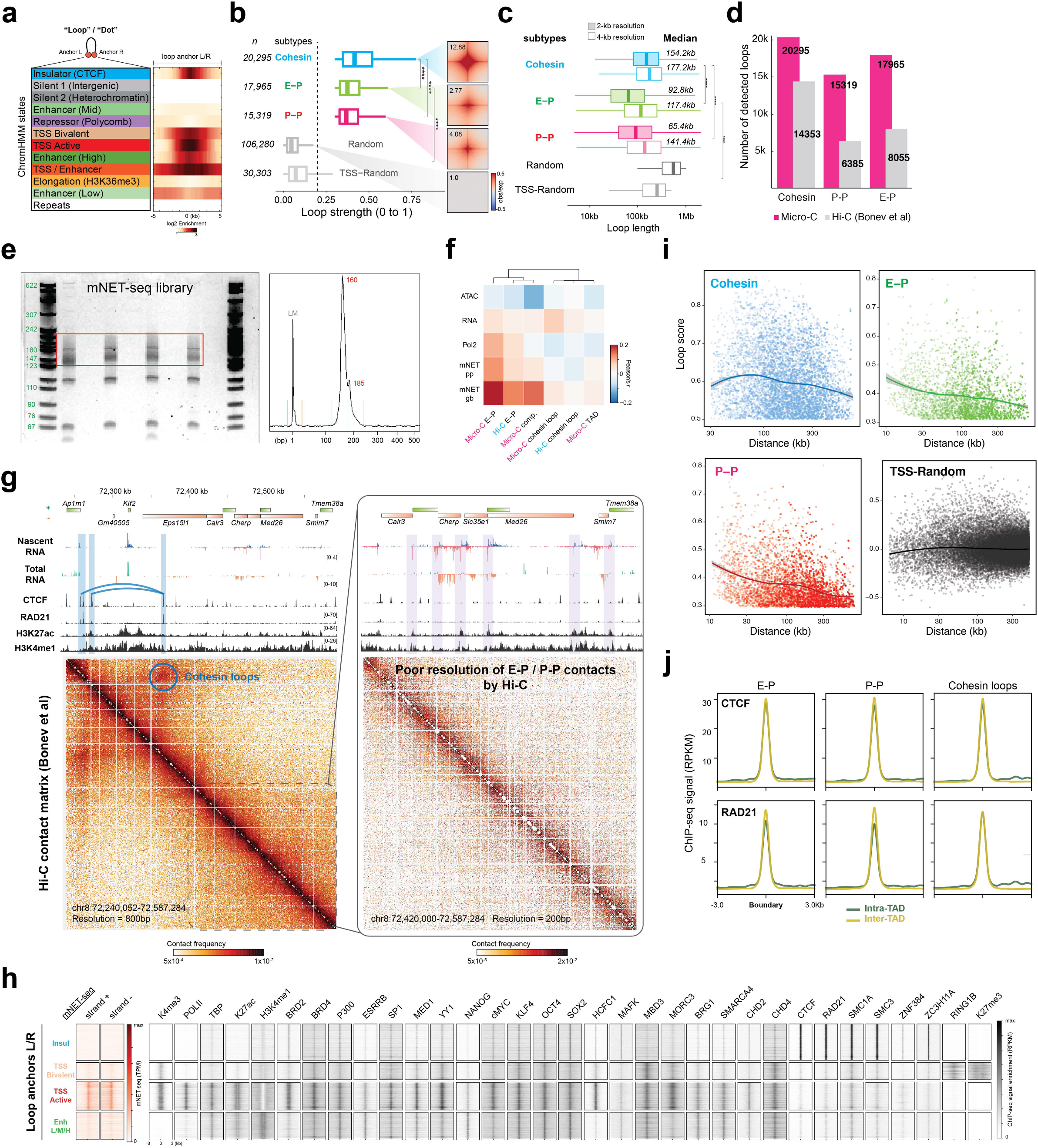
a. Enrichment of mESC ChromHMM^1^ states at loop anchors. The heatmap shows the log2 enrichment of each state in a ± 5-kb window around the loop anchors (loop number = 75,190 and loop anchor number = 118,733 after removing duplicates). b. Box plots of the loop strengths for cohesin, E-P, P-P, and random loops. Paired CTCF/cohesin, E-P, and P-P loci were quantified using Chromosight^2^ with Micro-C data at 2-kb or 4-kb resolution, which produced comparable but not identical results as Mustache^3^ in **Fig. 1b**. This loop list was used for the analysis in Extended Data **Fig. 1b-d**. The numbers of chromatin loops are shown on the left. The box plot indicates the quartiles for the loop strength score distribution. The genome-wide averaged contact signals (aggregate peak analysis (APA)) are plotted on the right. The contact map was normalized by matrix balancing and distance, with positive enrichment in red and negative signal in blue, shown as a diverging colormap with the gradient of normalized contact enrichment in log10. The ratio of contact enrichment for the center pixels is annotated within each plot. Asterisks denote a p-value < 10^−16^ by the Wilcoxon test. c. Box plots of the loop length distributions for cohesin, E-P, P-P, and random loops. The colored box represents the result using the 2-kb resolution Micro-C data and the white box using the 4-kb resolution data. The median size of loops is annotated on the right. We note that the median lengths are larger than our previous analysis with the insulation score due to the high computational expense to quantify the short-range loops with Micro-C data finer than 1-kb resolution. Asterisks denote a p-value < 10^−16^ by the Wilcoxon test. d. Summary of loop calling for cohesin, E-P, and P-P interactions with Micro-C (pink) or Hi-C data from Bonev et al.^4^ (gray). e. Gel image representing the size distribution of the mNET-seq library on a 6% PAGE gel and the resolved bands on the Fragment Analyzer electropherogram. f. Heatmap of Pearson’s correlation between the sequencing data (ATAC-seq, RNA-seq, Pol II ChIP-seq, and promoter pausing (pp) / gene body (gb) signal in mNET-seq) and the four levels of chromatin structures (compartment, TADs, cohesin loops, E-P and P-P loops) by Micro-C or Hi-C. g. Snapshots of Hi-C data in the same region as shown in **Fig. 1f**. h. Heatmaps of mNET-seq and ChIP-seq signal enrichments in a ± 3-kb window around the four primary types of loop anchors. i. Rank-ordered distribution of loop length against loop strength for cohesin, E-P, P-P, and random loops. The distribution for each loop type was fitted and smoothed by LOESS regression. The standard error (SE) is plotted as a gray shade along with the regression line. j. Profiles of ChIP-seq signal for CTCF and RAD21 at TAD boundaries grouped by intra-TAD (green) / inter-TAD (gold) cohesin, E-P, or P-P loops.

**Extended Data Fig 2.**
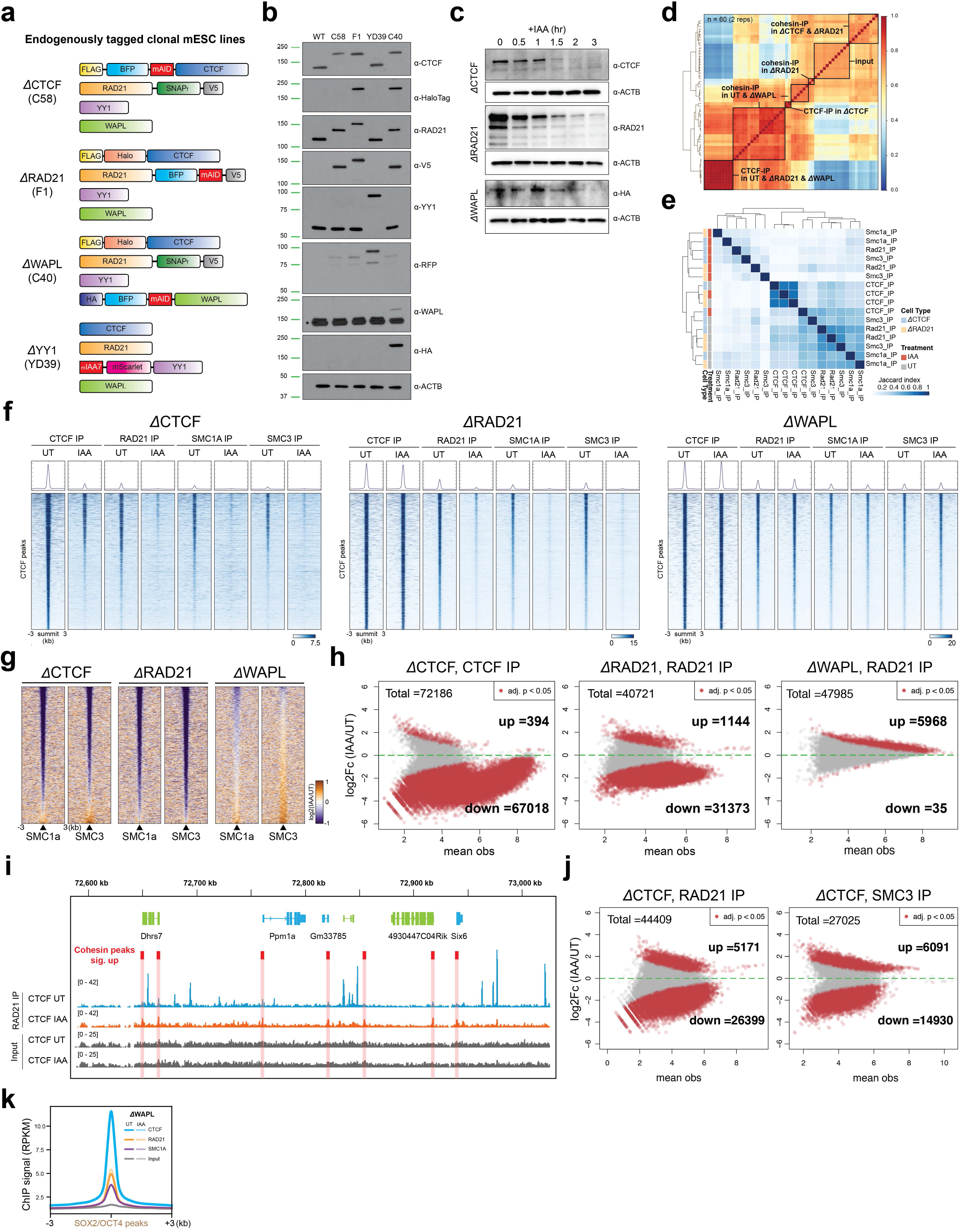
a. Schematics for endogenously tagging CTCF, RAD21, WAPL with the mAID degron and for endogenously tagging YY1 with miniIAA7. b. Immunoblots of CTCF, RAD21, WAPL, YY1, and their tags (HaloTag for CTCF, V5 for RAD21, RFP (mScarletI) for YY1, and HA for WAPL) for the protein expression levels and sizes in wildtype mESCs and in clones C58, F1, YD39, and C40. c. Immunoblots of CTCF, RAD21 and WAPL’s tag across a degradation time course from 0 (untreated) to 3 hr (IAA treatment) in clones C58, F1, and C40. d. Heatmap of ‘all-by-all’ Spearman’s correlation for all ChIP-seq replicates samples (n = 96). e. Heatmap of Jaccard’s index for the ratio of co-enriched peaks between the ChIP-seq replicates. f. Heatmaps of CTCF and cohesin (RAD21, SMC1A, and SMC3 subunits) ChIP-seq signal around WT-CTCF peaks called by MACS2 in the CTCF-, RAD21-, or WAPL-degron cells. The peaks called by MACS2 are plotted at the center across a ±3-kb region. The colormap shows the maximum signal (log2) in blue and the minimum signal in white. g. Heatmaps of differential ChIP-seq signals for SMC1A and SMC3 in cells depleted for CTCF, RAD21, or WAPL. The peaks called by MACS2 are plotted at the center across a ±3-kb region. The colormap shows an increased signal (log2) in orange and a decreased signal in purple after IAA treatment. h. MA plots show the differential ChIP-seq peaks between the UT and IAA-treated cells. The significantly changed peaks (adjusted *p*-value < 0.05) are colored in red. X-axis: mean observations of UT and IAA cells. Y-axis: log2 fold-change comparing the UT and IAA-treated cells. i. Snapshot of an example showing the position and level of cohesin at promoters (red shade) after CTCF degradation. j. MA plots showing the differential ChIP-seq peaks of RAD21 and SMC3 in the CTCF-depleted cells. The differential peaks (adjusted *p*-value < 0.05) are colored in red. X-axis: mean observations of UT and IAA cells. Y-axis: log2 fold-change comparing the UT and IAA-treated cells. k. Profiles of genome-wide averaged ChIP-seq for CTCF, RAD21, SMC1A, and input around the ±3-kb region of SOX2 or OCT4 peaks in WAPL-depleted cells.

**Extended Data Fig 3.**
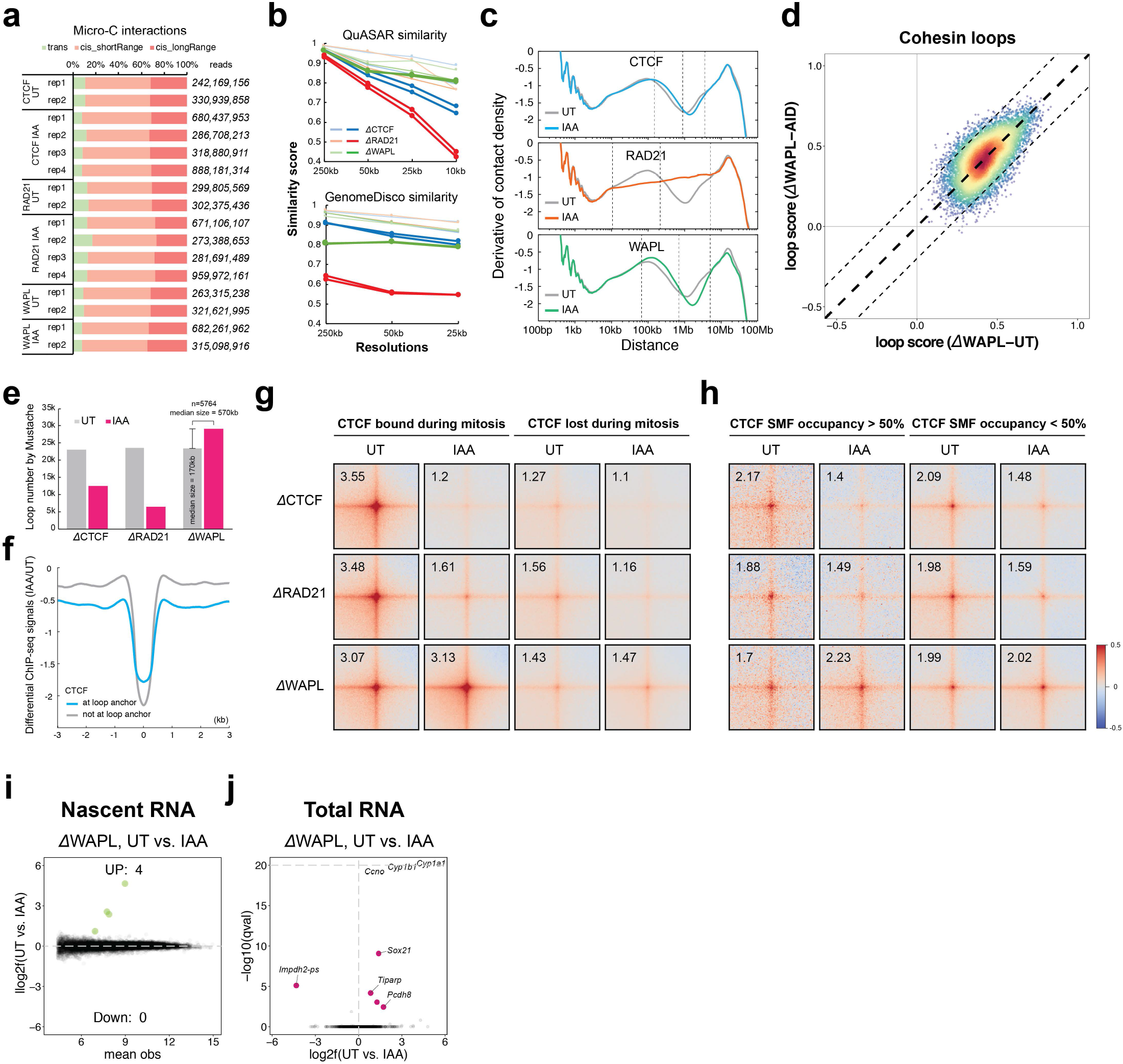
a. Summary of Micro-C experiments for CTCF, RAD21, and WAPL depletion. Total unique reads are annotated for each replicate on the right, consisting of trans-interactions (inter-chromosome) in green, short-range cis-interactions (< 20 kb) in orange, and long-range cis-interactions (> 20 kb) in red. b. Summary of reproducibility tests for Micro-C. Similarity scores measured by QuASAR (top panel) or GenomeDisco (bottom panel) either between replicates (light lines) or comparing the UT and IAA-treated samples (dark lines) using 250-kb, 50-kb, 25-kb, and 10-kb resolution of Micro-C matrices. c. Slope distribution of *P*(s) curves for the UT and IAA-treated cells. Dashed lines highlight the range of genome distances affected by CTCF, RAD21, or WAPL depletion. d. Scatter plot of loop scores for cohesin loops in the UT and IAA-treated cells. The overlaid heatmap indicates dot density, with the highest in red and the lowest in blue. Dashed lines along the diagonal mark the range that is characterized as unchanged loops. e. Summary of loop numbers called by Mustache for the UT (gray) and IAA-treated (pink) cells. The additional loops (n = 5764) identified after WAPL depletion show longer length with the median at 570 kb. f. Profile of differential ChIP-seq signal for CTCF in a ± 3-kb window around loop anchors (gray) or non-loop anchors (blue). g. APA for loops where CTCF is bound (left) or lost (right) at the anchors during mitosis^5^. h. APA for loops with either high (left) or low (right) CTCF occupancy characterized by single-molecular foot-printing (SMF) assay at their anchors^6^. i. MA plot of the mNET-seq result for WAPL depletion. DEGs with *q*-value < 0.05 are labeled with green (up) or pink (down). j. Volcano plot of total RNA-seq for WAPL depletion. DEGs (*q*-value < 0.01) are labeled with pink with the corresponding gene name.

**Extended Data Fig 4.**
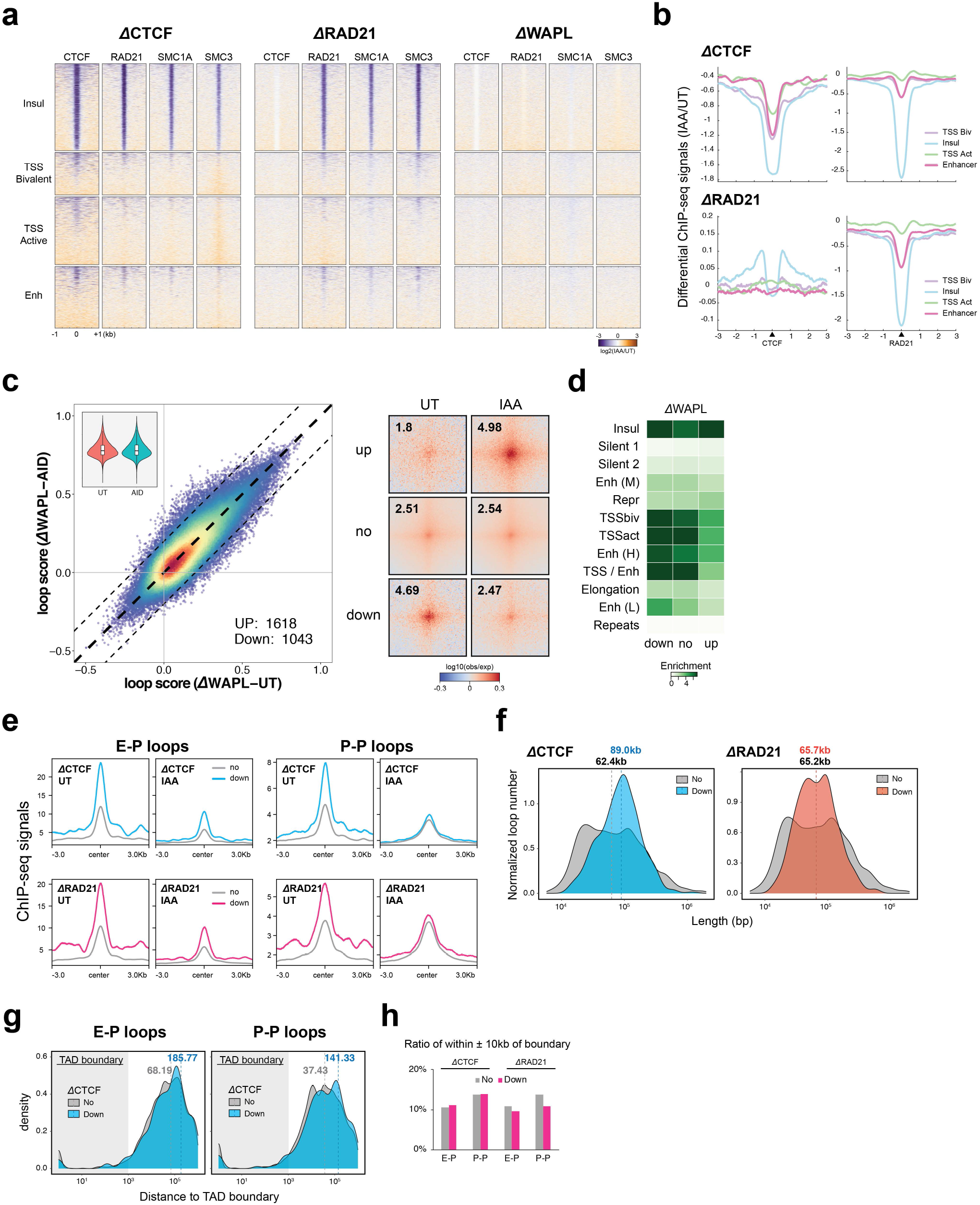
a. Heatmaps of differential ChIP-seq signals for CTCF, RAD21, SMC1A, and SMC3 comparing the UT and IAA-treated cells. Heatmaps were plotted across a ±3-kb region around four major types of loop anchors. The colormap shows an increased signal (log2) in orange and a decreased signal in purple after IAA treatment. b. Profiles of differential ChIP-seq signals for CTCF or RAD21 comparing the UT and IAA-treated cells across the same regions as in a. c. Scatter plot of loop scores for the called loops in the UT and IAA-treated cells (left). The violin chart (inset) shows the distribution of loop scores for the UT and IAA-treated conditions. The pile-up contact maps are plotted with loops sorted by up-regulated, down-regulated, or unchanged loops (right). d. Enrichment of the chromHMM states at loop anchors sorted by up-regulated, down-regulated, or unchanged after IAA treatment. e. Profiles of ChIP-seq signals for CTCF (top) or RAD21 (bottom) across a ±3-kb region around the anchors of E-P or P-P loops that are either unchanged (gray) or reduced after CTCF (blue) or RAD21 (pink) depletion. f. Histogram of the length distribution for loops that are unchanged (gray) or reduced in CTCF (blue) or RAD21 (orange) depletion. g. Length distribution of the unchanged or down-regulated E-P/P-P loops relative to TAD boundaries. h. Ratio of the unchanged (gray) or down-regulated (pink) E-P/P-P loop anchors located within ±10 kb of TAD boundaries.

**Extended Data Fig 5.**
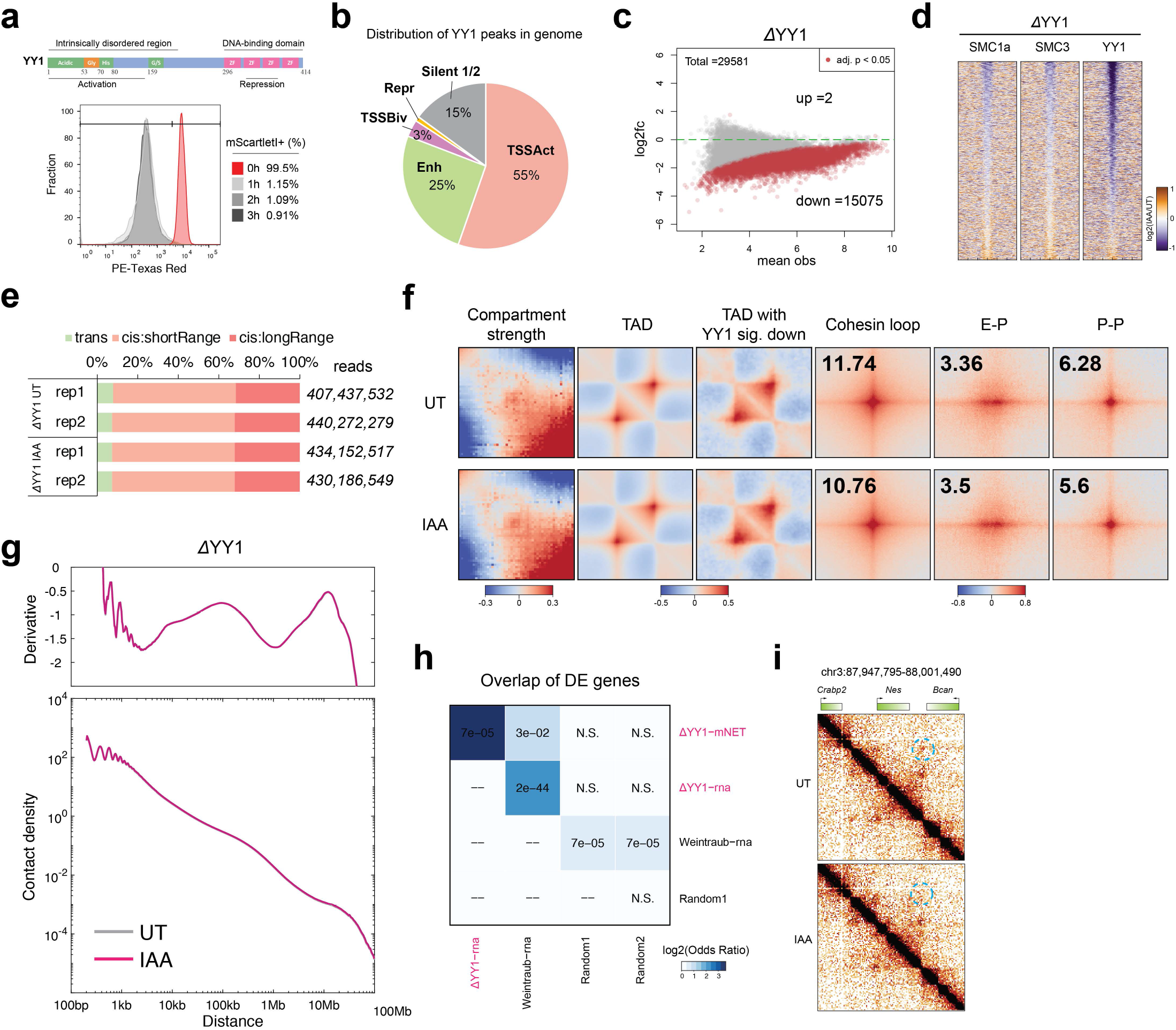
a. Schematic for YY1 protein domains (top) and histogram of mScarletI intensity for HaloTag-YY1 cells (clone YN11) treated with IAA for 0, 1, 2, or 3 hours. b. Pie chart shows the percentage of YY1 peaks enriched with four primary types of chromHMM states and silent chromatin. c. MA plots showing differential ChIP-seq peaks between the UT and IAA-treated cells. The significantly changed peaks (adjusted *p*-value < 0.05) are colored in red. X-axis: mean observations of the UT and IAA-treated cells. Y-axis: log2 fold-change comparing the UT and IAA-treated cells. d. Heatmaps of differential ChIP-seq signals for YY1, SMC1A, and SMC3 after YY1 depletion. e. Summary of Micro-C experiments for UT and YY1-depleted cells. Total unique reads are annotated for each replicate on the right, consisting of trans-interaction (inter-chromosome) in green, short-range cis-interaction (< 20 kb) in orange, and long-range cis-interaction (> 20 kb) in red. f. Genome-wide pileup plots for multiple levels of chromatin structures in the UT and IAA-treated cells. From left to right: 1) Saddle plot for compartmentalization strength. This shows average distance-normalized contact frequencies between 100-kb bins in cis with ascending eigenvector values (log2). Upper left and bottom right corners: contact frequency between B-B and A-A compartments. Upper right and bottom left corners: frequency of inter-compartment interactions. 2) Aggregate domain analysis (ADA) for TADs and TADs with significant YY1 ChIP-seq decrease. TADs were rescaled and aggregated at the center of the plot with matrix balancing normalization. 3) APA for cohesin, E-P, or P-P. The loops were aggregated at the center of a 20-kb window at 400-bp resolution. The genome-wide averaged loop enrichment was calculated by the fold enrichment (center pixel/expected upper-left pixels). g. Genome-wide contact decaying *P*(s) analysis (top) and the slope distributions of the *P*(s) curves for UT and IAA-treated cells. h. Overlap of DEGs between 1) mNET-seq after 3-hour YY1 depletion; 2) RNA-seq after 3-hour YY1 depletion; 3) RNA-seq after 3-hour YY1 depletion^7^; and 4) two sets of random genes. i. Snapshots of Micro-C maps comparing chromatin interactions in the UT (top) and IAA-treated (bottom) cells surrounding the *Nes* gene.

**Extended Data Fig 6.**
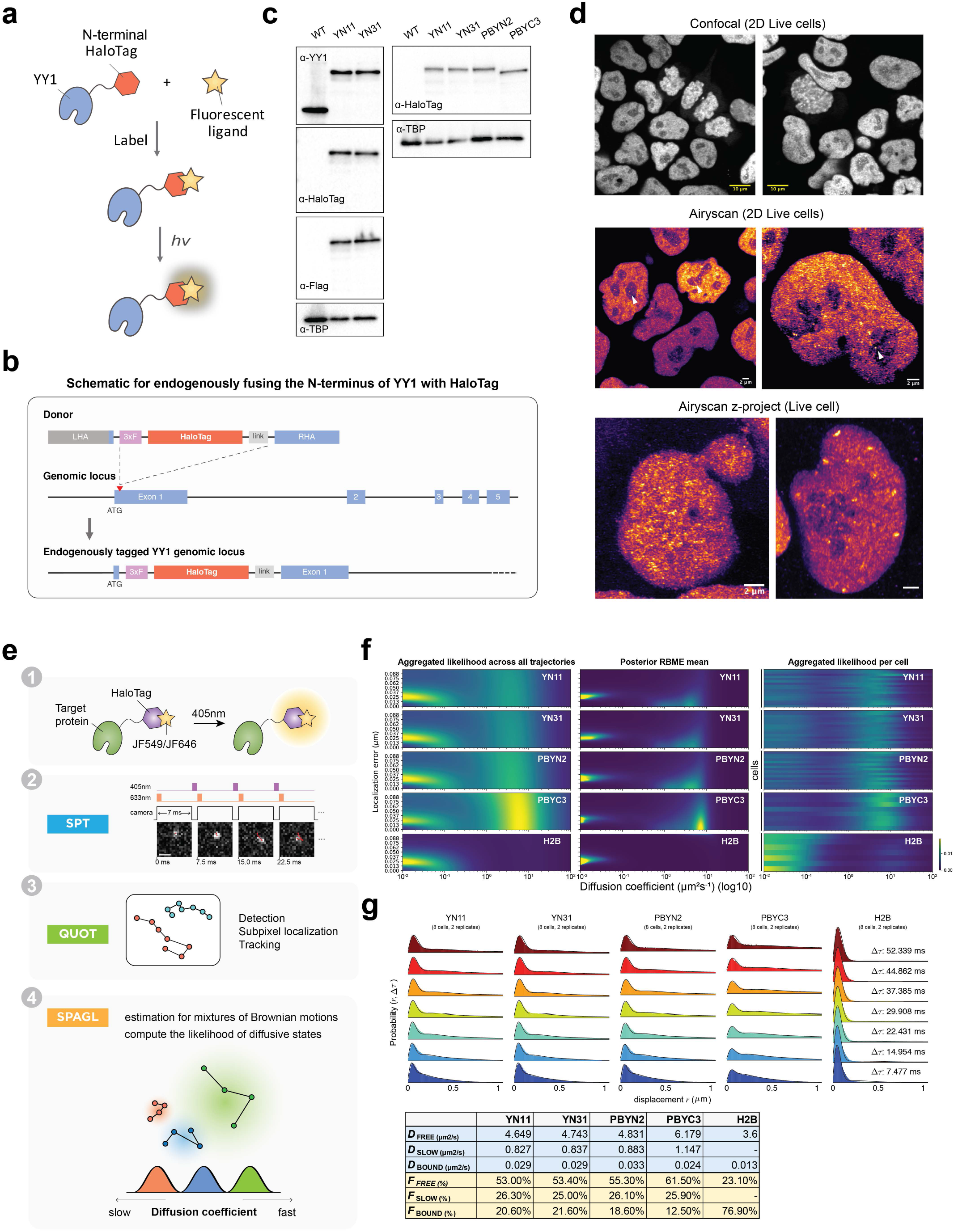
a. Schematic for conjugating a fluorescent dye with the HaloTag-YY1 fusion protein, which emits fluorescence upon excitation by a specific wavelength. b. Schematic for endogenously fusing the N-terminus of YY1 with HaloTag. c. Immunoblots of wild-type (WT), HaloTag-YY1 knock-in (YN11 and YN31), and stably expressing Halotag-YY1/YY1-HaloTag (PBYN2 and PBYC3) mESC lines for YY1, HaloTag, and FLAG proteins. TBP was used as a loading control. d. Confocal or Airyscan-resolved live-cell imaging for HaloTag-YY1 stained with 500 nM TMR. Arrow points to sporadic loci within the nucleolus. Images at the bottom panel are a z-projection with the mean signal. e. Schematic for the spaSPT experiment and the analysis pipeline with Quot and Spagl^8^. f. Heatmaps of localization errors obtained by aggregated likelihood across all trajectories (left) or posterior marginalized localization error (middle) for clone YN11, YN31, PBYN2, PBYC3 and H2B. The distribution of the likelihood of diffusion coefficients (x-axis) for single cells (each row at y-axis) is plotted on the right panel. g. spaSPT displacement histograms for YN11, YN31, PBYN2, PBYC3, and H2B. Raw displacement data for seven different lag times are shown with a three-state Spot-On model^9^ fit overlaid. The inferred fractions and diffusion coefficients for each cell are shown in the table in the bottom panel.

**Extended Data Fig 7.**
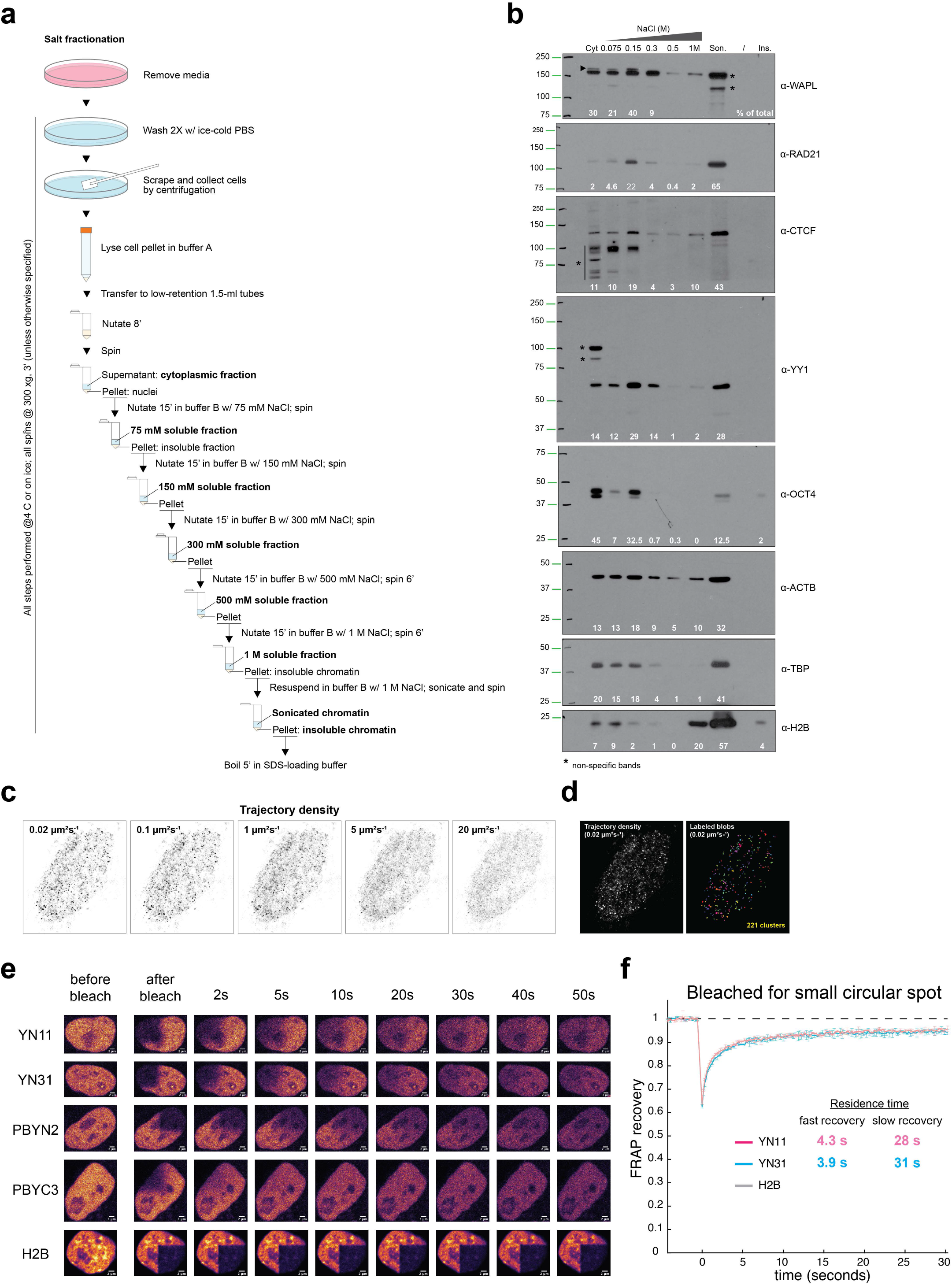
a. Schematic for biochemical fractionation experiment. b. Immunoblots of cytoplasmic (Cyt) and nuclear proteins dissociating from chromatin at increasing salt concentrations (75, 150, 300, 500 mM and 1M NaCl) as schematized in a, probed with the indicated antibodies (*α*). Son. Sonicated chromatin; Ins. Insoluble pellet after chromatin sonication; % of total: signal intensity of each fraction divided by the total signal intensity across all fractions. In the top blot, the arrowhead indicates the band corresponding to WAPL; asterisks denote non-specific bands. c. Spatial reconstruction of spaSPT data for YY1’s trajectory densities. YY1 trajectories are binned by diffusion coefficients, as indicated. d. Segmentation of YY1 clusters with the reconstructed images from the spaSPT data. Individual YY1 clusters are colored. e. Snapshots of FRAP experiments for multiple time points from “before bleach” to “50 sec after bleach”. f. FRAP analysis of YY1 bleached with a small circular spot. Error bars indicate the standard deviation of each acquired data point.

**Extended Data Fig 8.**
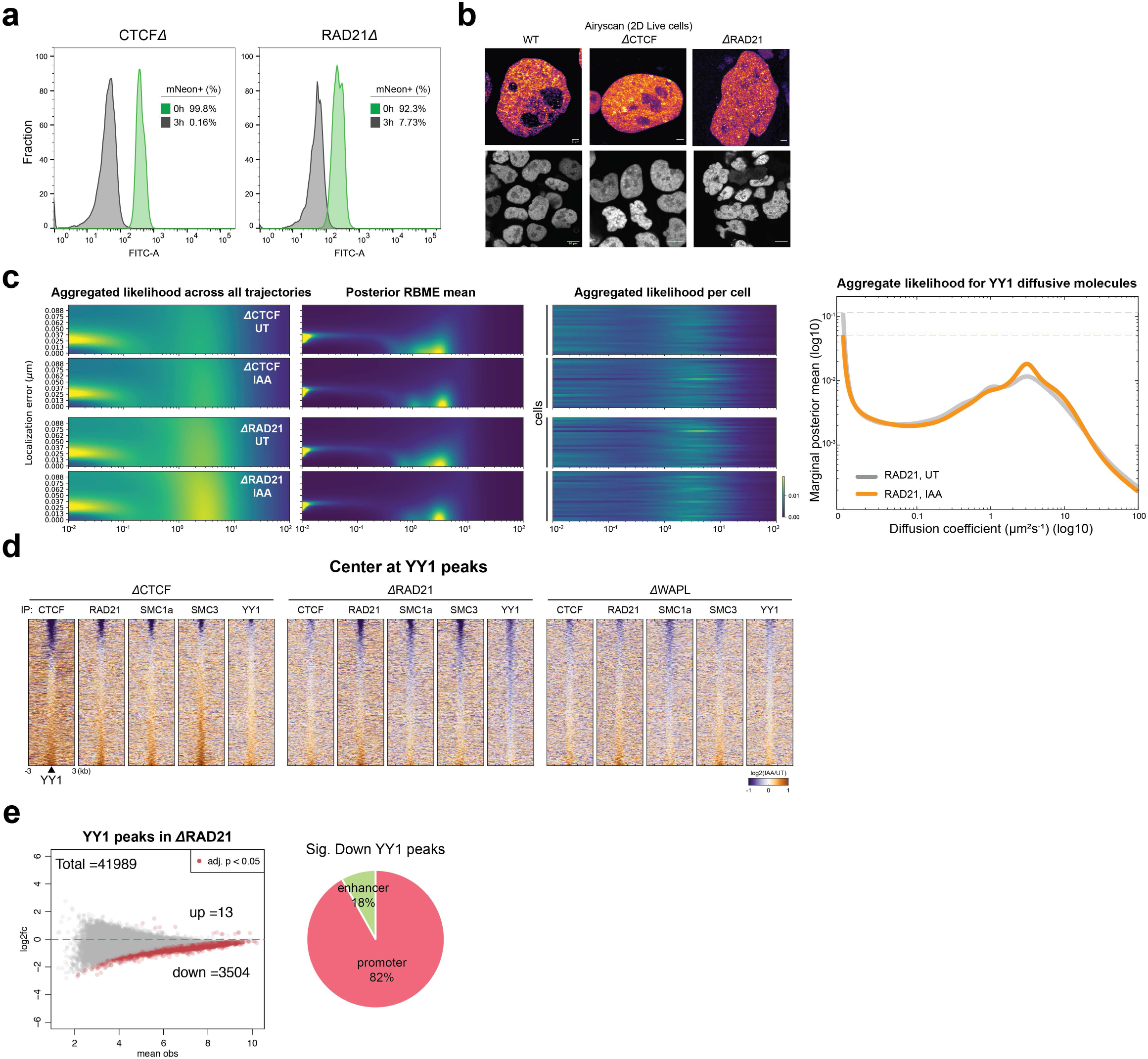
a. Histogram of mNeonGreen intensity for cells (clones CD1 and RD35) treated with IAA for 0 or 3 hours. b. Airyscan-resolved live-cell imaging for HaloTag-YY1 stained with 500 nM TMR in wild type, CTCF-, or RAD21-depleted cells. c. Heatmaps of localization errors obtained by aggregated likelihood across all trajectories (panel one from left) or posterior marginalized localization error (panel two) for the UT or IAA-treated cells. The distribution of likelihood of diffusion coefficients (x-axis) for single cells (each y-axis row) is plotted on panel three. Estimation of YY1 diffusion coefficients by regular Brownian motion with marginalized localization errors is plotted on panel four. d. Heatmaps of differential ChIP-seq signals for CTCF, RAD21, SMC1a, SMC3, and YY1 in cells depleted for CTCF, RAD21, and WAPL. The peaks are centered on wildtype YY1 peaks across a ±3-kb region. The colormap shows an increased signal (log2) in orange and a decreased signal in purple after IAA treatment. e. MA plot showing the differential ChIP-seq peaks between the UT and IAA-treated cells (left). The significantly changed peaks (adjusted *p*-value < 0.05) are colored in red. X-axis: mean observations of UT and IAA cells. Y-axis: log2 fold-change comparing the UT and IAA-treated cells. Pie chart shows the percentages of the down-regulated YY1 peaks enriched at promoters and enhancers (right).

## Methods

### Cell culture, stable cell line construction and dye labeling

JM8.N4 mouse embryonic stem cells (mESCs)^1^ (Research Resource Identifier: RRID:CVCL_J962; obtained from the KOMP Repository at UC Davis) were used for all experiments. Cells were cultured on plates pre-coated with 0.1% gelatin (Sigma-Aldrich G9291) in knock-out DMEM (ThermoFisher #10829018) supplemented with 15% FBS (HyClone FBS SH30910.03 lot #AXJ47554), 0.1 mM MEM non-essential amino acids (ThermoFisher #11140050), 2 mM GlutaMAX (ThermoFisher #35050061), 0.1 mM 2-mercaptoethanol (Sigma-Aldrich M3148), 1% Penicillin-Streptomycin (ThermoFisher #15140122), and 1000 units of LIF (Millipore). Media was replaced daily and cells were passaged every 2 days by trypsinization. Cells were grown at 37°C and 5.5% CO2 in a Sanyo copper alloy IncuSafe humidified incubator (MCO-18AIC(UV)). For imaging, the media was identical except that knock-out DMEM lacking phenol red (ThermoFisher #31053028) was used to minimize background fluorescence.

Cell lines stably expressing 3xFLAG-HaloTag-YY1 and YY1-HaloTag-3xFLAG were generated using PiggyBac transposition and drug selection. The coding sequences of YY1 and 3xFLAG-HaloTag or HaloTag-3xFLAG were cloned by Gibson Assembly into a PiggyBac vector co-expressing a puromycin resistance gene. For each construct, YY1 and HaloTag were connected by a short peptide linker sequence (GDGAGLING). 1.3 µg of PiggyBac vector was co-transfected with 0.5 µg of SuperPiggyBac transposase vector into JM8.N4 cells by nucleofection using the Lonza Mouse Embryonic Stem Cell Nucleofector Kit reagents and the Amaxa Nucleofector II device. After 24 hr, antibiotic selection was begun with 500 ng/mL of puromycin. Once cells reached ∼80% confluence, 5000 cells were seeded onto a 15 cm plate and allowed to grow into colonies. After 4-6 days, single clones were isolated and seeded onto a 96-well plate. Cells were allowed to expand for 4-5 days, and successfully integrated PiggyBac constructs were validated by PCR genotyping. We selected clone PBYN2 for 3xFLAG-HaloTag-YY1 and clone PBYC3 for YY1-HaloTag-3xFLAG.

For all single-molecule experiments, cells were grown overnight on Matrigel-coated 25-mm circular no 1.5H cover glasses (High-Precision 0117650). Prior to all experiments, the cover glasses were plasma-cleaned and then stored in isopropanol until use. Cells were labeled with 50 nM PA-JFX646 (fast SPT) or 25 pM JFX549 (slow SPT) HaloTag ligand for 30 min and washed twice with PBS. After the final washes, cells were replenished with phenol-free medium prior to imaging.

For PALM experiments, cells were grown overnight on Matrigel-coated 25-mm circular no 1.5H cover glasses (High-Precision 0117650). Prior to all experiments, the cover glasses were plasma-cleaned and then stored in isopropanol until use. Cells were labeled with 500 nM PA-JFX549 HaloTag ligand for 30 min, washed twice with fresh media for 5 min, and then washed once with pH 7.4 phosphate-buffered saline (PBS). Labeled cells were fixed with 4% paraformaldehyde and 2% glutaraldehyde in PBS for 20 min at 37°C, washed once with PBS, and imaged in PBS with 0.01% (w/v) NaN_3_.

For FRAP experiments, cells were grown overnight on Matrigel-coated (Corning #354277; purchased from ThermoFisher #08-774-552) glass-bottom 35-mm dishes (MatTek P35G-1.5-14C). Cells were labeled with 500 nM TMR for 30 min and washed twice with PBS.

### Generation of CRISPR/Cas9-mediated knock-in cell lines

Endogenously tagged mESC lines were generated by CRISPR/Cas9-mediated genome editing as previously described^2^ with modifications. mESCs were seeded to 6-well plates and each well was transfected with Lipofectamine 3000 (ThermoFisher #L3000015) or nucleofection (Lonza #VPH-1001) using 1 µg of a construct encoding Cas9, Venus yellow fluorescent protein, and a sgRNA and 2 µg of a homology repair plasmid with the DNA insert of interest flanked by left and right homology arms (∼500 bp each). For each insertion we designed 2-4 individual sgRNAs with the CRISPOR tool^3^ to be transfected separately into mESCs and pooled before sorting Venus-positive cells by FACS. Sorted cells were plated at a density of ∼5,000-10,000 cells per P15 plate and media replaced every other day. Single colonies were transferred to 96-well plates to be expanded and genotyped by PCR after crude DNA extraction with the DirectPCR Lysis Reagent (Viagen Biotech #302-C). Clones with a correct insert were further verified by Sanger sequencing and Western blot.

The parental cell line of CTCF, Rad21 and WAPL auxin-inducible degron (AID) mESCs was clone C59 from Hansen et al.^4^ further modified to visualize DNA looping (Clone C36 osTir1: full details on the generation of C36 will be published elsewhere (in preparation) and are available upon request).

To endogenously tag 1) CTCF, RAD21, or WAPL with mAID^5^ in the C36 osTir mESC line; 2) YY1 with miniIAA7^6^ in the parental JM8.N4 with osTIR1 expression; 3) YY1 with HaloTag in the parental JM8.N4; and 4) CTCF and RAD21 with mAID in the YY1-HaloTag cell lines, we designed homology repair plasmids encoding 1) 3xFLAG-BFP-mAID-CTCF, RAD21-BFP-mAID-V5, HA-BFP-mAID-WAPL; 2) miniIAA7-mScarletI-YY1; 3) 3xFLAG-HaloTag-YY1; 4) 3xFLAG-mNeonGreen-mAID-CTCF and RAD21-mNeonGreen-mAID-V5 (plasmids, sgRNAs and primer sequences in **Supplementary Table 1**). Homozygous clones with the correct genotype were further verified by Sanger sequencing and Western blot after IAA-mediated depletion. Among the several clones generated for each AID line, we picked clone C58 for 3xFLAG-BFP-mAID-CTCF; clone F1 for RAD21-BFP-mAID-V5; clone C40 for HA-BFP-mAID-WAPL; clone YD39 for miniIAA7-mScarletI-YY1; clone YN11 for 3xFLAG-HaloTag-YY1; clone CD1 for 3xFLAG-mNeonGreen-mAID-CTCF and clone RD35 for RAD21-mNeonGreen-mAID-V5.

### Western Blotting

For Western blot analysis, cells were seeded to 6-well plates, washed with ice-cold PBS and scraped in 300 μl of low-salt lysis buffer (0.1 M NaCl, 25 mM HEPES, 1 mM MgCl_2_, 0.2 mM EDTA, 0.5% NP-40 and protease inhibitors), with 125 U/mL of benzonase (Novagen, EMD Millipore), rocked at 4 °C for 1 hr and then NaCl was added to a 0.2-M final concentration. Lysates were then rocked at 4 °C for 30 min and centrifuged at maximum speed at 4 °C. Supernatants were quantified by Bradford. Alternatively, cells were lysed directly on plates with 300 μl of high-salt lysis buffer (0.5 M NaCl, 25 mM HEPES, 1 mM MgCl_2_, 0.2 mM EDTA, 0.5% NP-40 and protease inhibitors) and transferred to low-stick tubes with 100 µl of 4x SDS-loading buffer. Proteins were loaded onto a 9% Bis-Tris SDS-PAGE gel, transferred onto a nitrocellulose membrane (Amershan Protran 0.45 um NC, GE Healthcare) for 1 hr at 100V, blocked in TBS-Tween with 10% milk for at least 1 hr at room temperature and blotted overnight at 4 °C with primary antibodies in TBS-T with 5% milk (antibodies listed in **Supplementary Table 1**). HRP-conjugated secondary antibodies were diluted 1:5000 in TBS-T with 5% milk and incubated with the membrane at room temperature for an hour before performing the chemiluminescence reaction (Western Lightning Plus-ECL, Enhanced Chemiluminescence Substrate, Perkin Elmer NEL105001EA). Signal was captured with either X-ray films (CL-XPosure™ Film, ThermoScientific 34091) or with a Bio-Rad ChemiDoc imaging system.

### Chromatin immunoprecipitation (ChIP)

ChIP was performed as described with a few modifications^7^. Cells were treated with either ethanol or 500 mM of ethanol-dissolved auxin for 3 hrs and cross-linked for 7 min at room temperature with 1% formaldehyde-containing medium; cross-linking was stopped by PBS-glycine (0.125 M final). Cells were washed twice with ice-cold PBS, scraped, centrifuged for 10 min and pellets were flash-frozen. Cell pellets were thawed and resuspended in cell lysis buffer (5 mM PIPES, pH 8.0, 85 mM KCl, and 0.5% NP-40, 1 ml/15 cm plate) with protease inhibitors and incubated for 10 min on ice. Lysates were centrifuged for 10 min at 1250 rcf and nuclear pellets were resuspended in 6 volumes of sonication buffer (50 mM Tris-HCl, pH 8.1, 10 mM EDTA, 0.1% SDS) with protease inhibitors, incubated on ice for 10 min, and sonicated to obtain DNA fragments below 2000 bp in length (Covaris S220 sonicator, 20% Duty factor, 200 cycles/burst, 150 peak incident power, 10-20 cycles 50 sec on and 30 sec off). Sonicated lysates were cleared by centrifugation and chromatin (400 μg per antibody) was diluted in RIPA buffer (10 mM Tris-HCl, pH 8.0, 1 mM EDTA, 0.5 mM EGTA, 1% Triton X-100, 0.1% SDS, 0.1% Na-deoxycholate, 140 mM NaCl) with protease inhibitors to a final concentration of 0.8 μg/μl, precleared with Protein G sepharose (GE Healthcare) for 2 hrs at 4°C and immunoprecipitated overnight with 4 μg of specific antibodies (antibodies listed in **Supplementary Table 1**). About 4% of the precleared chromatin was saved as input. Immunoprecipitated DNA was purified with the Qiagen QIAquick PCR Purification Kit, eluted in 36 μl of 0.1X TE (1 mM Tris-HCl pH 8.0, 0.01 mM EDTA) and analyzed by qPCR together with 2% of the input chromatin prior to ChIP-seq library preparation (SYBR® Select Master Mix for CFX, ThermoFisher; ChIP-qPCR primer sequences in **Supplementary Table 1**).

ChIP-seq libraries were prepared using the NEBNext^®^ Ultra^™^ II DNA Library Prep Kit for Illumina^®^ (NEB E7645) according to manufacturer instructions with a few modifications. 20 ng of ChIP input DNA (as measured by Nanodrop) and 25 μl of the immunoprecipitated DNA were used as a starting material and the recommended reagent volumes were cut in half. The NEBNext Adaptor for Illumina was diluted 1:10 in Tris/NaCl, pH 8.0 (10 mM Tris-HCl pH 8.0, 10 mM NaCl) and the ligation step extended to 30 min. After ligation, a single purification step with 0.9X volumes of Agencourt AMPure XP PCR purification beads (Beckman Coulter A63880) was performed, eluting DNA in 22 μl of 10 mM Tris-HCl pH 8.0. 20 μl of the eluted DNA was used for the library enrichment step, performed with the KAPA HotStart PCR kit (Roche Diagnostics KK2502) in 50 μl of total reaction volume (10 μl 5X KAPA buffer, 1.5 μl 10 mM dNTPs, 0.5 μl 10 μM NEB Universal PCR primer, 0.5 μl 10 μM NEB index primer, 1 μl KAPA polymerase, 16.5 μl nuclease-free water and 20 μl sample). Samples were enriched with 9 PCR cycles (98 °C, 45 sec; [98 °C, 15 sec; 60 °C, 10 sec] x 9; 72 °C, 1 min; 4 °C, hold), purified with 0.9 volumes of AMPure XP PCR purification beads and eluted with 33 μl of 10 mM Tris-HCl pH 8.0. Library concentration, quality and fragment size were assessed by Qubit fluorometric quantification (Qubit™ dsDNA HS Assay Kit, Invitrogen^TM^ Q32851), qPCR and Fragment analyzer™. 12 multiplexed libraries were pooled and sequenced in one lane on the Illumina HiSeq4000 sequencing platform (50-bp, single end-reads) at the Vincent J. Coates Genomics Sequencing Laboratory at UC Berkeley, supported by NIH S10 OD018174 Instrumentation Grant.

ChIP-seq raw reads from ethanol (UT)- or auxin (IAA)-treated ΔCTCF, ΔRAD21, ΔWAPL and ΔYY1 mESCs (96 libraries total, 2 biological replicates per condition) were quality-checked with FastQC and aligned onto the mouse genome (mm10 assembly) using Bowtie^8^, allowing for two mismatches (-n 2) and no multiple alignments (-m 1). Biological replicates were pooled and peaks were called with MACS2 (`-- nomodel --extsize 250`)^9^ using input DNA as a control. To create heatmaps we used deepTools (version 2.4.1)^10^. We first ran bamCoverage (`--binSize 50 --normalizeTo1x 2150570000 - -extendReads 300 --ignoreDuplicates -of bigwig`) and normalized read numbers to 1x sequencing depth, obtaining read coverage per 50-bp bins across the whole genome (bigWig files). We then used the bigWig files to compute read numbers across 6 kb centered on either CTCF, RAD21, or YY1 peak summits as called by MACS2 in UT cells (`computeMatrix reference-point -- referencePoint=TSS --upstream 3000 --downstream 3000 –missingDataAsZero --sortRegions=no`). We sorted the output matrices by decreasing UT enrichment, calculated as the total number of reads within a MACS2 called ChIP-seq peak. Finally, heatmaps were created with the plotHeatmap tool (`--averageTypeSummaryPlot=mean --colorMap=’Blues’ -- sortRegions=no`). To identify the differential peaks between UT- and IAA-treated cells, we used MAnorm2^11^, which employs a hierarchical strategy for normalization of ChIP-seq data and assesses within-group variability of ChIP-seq signals based on an empirical Bayes framework. We note that the total ChIP-seq peak numbers in MAnorm2 are combined from UT- and IAA-treated cells and may differ from the number of MACS2 calling.

### Biochemical fractionation

Wild type JM8.N4 mESCs were seeded to 15-cm plates, washed with ice-cold PBS, scraped in PBS and pelleted at 135 rcf, 10 min at 4 °C. Pellets were resuspended in 350 μl of cell lysis buffer A (10 mM HEPES pH 7.9, 10 mM KCl, 3 mM MgCl_2_, 340 mM Sucrose, 10% glycerol v/v, 1 mM DTT and freshly added 0.1% Triton X-100 v/v and protease inhibitors) and rocked for 8 min at 4 °C. Nuclei were pelleted at 3,000 xg for 3 min at 4 °C and the supernatant containing the cytoplasmic fraction was saved. Nuclei were resuspended in 350 μl of buffer B with 75 mM NaCl (9 mM EDTA, 0.2 mM EGTA, 1 mM DTT and freshly added 0.1% Triton X-100 v/v and protease inhibitors) and rocked at 4 °C for 15 min. Nuclei were pelleted again as above (supernatant saved as the 75 mM wash fraction) and washed with 350 μl of buffer B with increasing NaCl concentrations (150 mM, 300 mM, 500 mM and 1 M, see **Extended Data Figure 7** for a step-by-step procedure). After collecting the 1 M wash, the pellet was resuspended to 350 μl of 1 M buffer B and sonicated (Covaris S220 sonicator, 20% Duty factor, 200 cycles/burst, 100 peak incident power, 8 cycles 20 sec on and 40 sec off). The sonicated lysate was spun down and the insoluble pellet boiled in SDS-loading buffer. 10 μl of each fraction was added to 2 μl of 4X SDS loading buffer and subjected to Western blot as detailed above. Band intensities were quantified with the ImageJ “Analyze Gels” function^12^.

### Micro-C assay for mammalian cells

We briefly summarize the Micro-C experiment here. The detailed protocol and technical discussion are available in our previous study^13^. Mouse embryonic stem cells (JM8.N4) and the derivative genome-edited lines were cultured in the recommended conditions^1^. When cells grew to ∼70% confluency, we resuspended them in 0.05% of trypsin, inactivated with cell culture media, and resuspended in 1% formaldehyde crosslinking media (without FBS). Cells were crosslinked for 10 min at room temperature (RT) while nutating. We then added 1 M Tris-HCl pH 7.5 (final concentration 375 mM) and incubated for 5 min at RT to quench the crosslinking reaction. Cells were spun down and washed twice with cold PBS. Cell pellets were crosslinked again with the freshly prepared 3 mM DSG crosslinking solution (in base media without FBS) for 45 min at RT while nutating. The crosslinking reaction was quenched and washed following the same steps as above. We routinely split cells into 1 million cells per vial after fixation and perform MNase titration and Micro-C with 1 million cells. Crosslinked cell pellets can be snap frozen in liquid nitrogen and stored at −80 °C for months or used immediately for the next step. We note that 1) cells directly crosslinked on the dish typically yield a similar result; 2) using freshly made formaldehyde and DSG solution is critical to obtain high-quality Micro-C data; and 3) to avoid loss of cells, low-retention tubes and tips are strongly recommended.

Crosslinked cell pellets were resuspended in MB1 (50 mM NaCl, 10 mM Tris-HCl pH 7.5, 5 mM MgCl_2_, 1 mM CaCl_2_, 0.2% NP-40) at a concentration of 1×10^6^ cells/100 µL and were incubated for 20 min on ice. Cells were spun down and washed once with MB1. We then added the appropriate amount of Micrococcal nuclease (MNase) and incubated the tube for 10 min at 37 °C while shaking in a thermomixer at ∼850 rpm. The optimal digestion condition results in ∼90% of mono-nucleosome and ∼10% of di-nucleosome. The MNase reaction was inactivated by adding 4 mM EGTA and incubated for 10 min at 65 °C. Digested chromatin was spun down and washed twice with 1 mL of cold MB2 (50 mM NaCl, 10 mM Tris-HCl pH 7.5, 10 mM MgCl_2_). We note that and MNase titration that yields 90% monomer/10% dimers substantially reduces contamination with un-digested (un-ligated) dimers in Micro-C data.

MNased-fragmented chromatin was then subjected to the three-step end-repair protocol to generate ligatable ends filled with biotin-dNTPs. First, the pellet was resuspended in the end-repair buffer (50 mM NaCl, 10 mM Tris-HCl pH 7.5, 10 mM MgCl_2_, 100 µg/mL BSA, 2 mM ATP, 5 mM DTT) and the 5’-ends were phosphorylated with 25 units of T4 Polynucleotide Kinase (NEB #M0201) for 15 min at 37 °C while shaking in a thermomixer at 1000 rpm for an interval of 15 sec every 3 min. Second, to convert the mixed types of nucleosomal ends to cohesive ends, we added 25 units of DNA Polymerase I, Large (Klenow) Fragment (NEB #M0210) directly to the reaction and incubated the tube for an additional 15 min at 37 °C while shaking in a thermomixer at 1000 rpm for an interval of 15 sec every 3 min. Third, to repair the nucleosomal DNA ends to the blunt and ligatable ends, we supplemented 66 µM of dNTPs (dTTP, dGTP (NEB #N0446), biotin-dATP (Jena Bioscience #NU-835-BIO14), biotin-dCTP (Jena Bioscience #NU-809-BIOX), and 1X T4 DNA ligase reaction buffer (NEB #B0202) directly into the reaction and incubated for 45 min at 25 °C while shaking in a thermomixer at 1000 rpm for an interval of 15 sec every 3 min. The end-repair reaction was then inactivated with 30 mM EDTA for 20 min at 65°C without shaking. Next, chromatin was pelleted by centrifugation for 5 min at ∼10,000xg at 4 °C and washed once with cold MB3 (50 mM Tris-HCl pH 7.5, 10 mM MgCl_2_).

End-repaired nucleosomes were then subjected to proximity ligation with ∼5000 cohesive end units (CEU) of T4 DNA ligase (NEB #M0202) in 1X T4 DNA ligase reaction buffer (NEB #B0202) for at least 2 hrs at room temperature with slow rotation on an orbital shaker. Biotin-dNTPs at the unligated DNA termini were removed by ∼500 units of Exonuclease III (NEB #M0206) in 1X NEBuffer 1 (NEB #B7001) for 15 min at 37°C while shaking at 1000 rpm for an interval of 15 sec every 3 min. Samples were then reverse crosslinked with 1X proteinase K solution (500 ug/uL Proteinase K (ThermoFisher #AM2542), 1% SDS, 0.1 µg/µL RNaseA) at 65°C overnight. DNA was extracted by the standard phenol:chloroform:isoamyl alcohol (25:24:1) and ethanol precipitation procedure. DNA was then purified again with the ZymoClean DNA Clean & Concentrator-5 Kit (Zymo #D4013). Purified DNA was separated on a 3% NuSieve GTG agarose gel (Lonza #50081). The gel band corresponding to the size of dinucleosomal DNA (∼250 to 350 bp) was cut and purified with the Zymoclean Gel DNA Recovery Kit (Zymo #D4008). We note that size selection for DNA larger than 200 bp greatly reduces the ratio of unligated monomers in Micro-C data.

Micro-C sequencing libraries were generated by using the NEBNext Ultra II DNA Library Prep Kit for Illumina (NEB #E7645) with some minor modifications. We first repaired the purified DNA again using the End-it DNA End-Repair Kit (Lucigen #ER0720) following the manufacturer’s suggested conditions. The mix was incubated for 45 min at 25 °C and then inactivated the enzyme reaction for 10 min at 70 °C. This step is optional, but we find it increases the library yield. Biotinylated DNA was captured by Dynabeads MyOne Streptavidin C1 beads (ThermoFisher #65001) in 1X BW buffer (5 mM Tris-HCl pH 7.5, 0.5 mM EDTA, 1 M NaCl) on a nutator for 20 min at room temperature. Beads were washed twice with 1X TBW buffer (0.1% Tween20, 5 mM Tris-HCl pH 7.5, 0.5 mM EDTA, 1 M NaCl) for 5 min at 55 °C while shaking at 1200 rpm, rinsed once with Tris buffer (10 mM Tris-HCl pH 7.5), and then resuspended in Tris buffer.

We then performed ‘on-bead’ end-repair/A tailing and adapter ligation following the NEB protocol. After adapter ligation, beads were washed once with 1X TBW and rinsed once with Tris buffer. The Micro-C library was generated by using the KAPA HiFi HotStart ReadyMix (Roche #KK2601) or the Q5 High-Fidelity 2X Master Mix (NEB #E7645) with the manufacturer’s suggested conditions. We recommend using a minimal PCR cycle to reduce PCR duplicates, typically 8 to 12 cycles, which can generate high-quality Micro-C data. Prior to sequencing, purifying the library twice with 0.85X AMPure XP beads (Beckman #A63880) can eliminate primer dimers and adapters. We used Illumina 100 bp paired-end sequencing (PE100) to obtain ∼400M reads for each replicate in this study.

### Micro-C data processing and analyses

Valid Micro-C contact read pairs were obtained from the HiC-Pro analysis pipeline^14^, and the detailed description and code can be found at https://github.com/nservant/HiC-Pro. In brief, paired fastq files were mapped to the mouse mm10 genome independently using Bowtie 2.3.0 with ‘very-sensitive-local’ mode^8^. Aligned reads were paired by the read names. Pairs with multiple hits, low MAPQ, singleton, dangling end, self-circle, and PCR duplicates were discarded. Paired reads with distances shorter than 100 bp (e.g., unligated mono-nucleosome) were also removed. Output files containing all valid pairs were used in downstream analyses. We recommend running a pilot sequencing run (∼10M reads) and checking the following quality control statistics before moving forward to a high-coverage sequencing: (1) bowtie mapping rate; (2) reads pairing percentage; (3) ratio of sequencing artifacts; (4) ratio of cis/trans contacts; (5) unligated monomer percentage. If any of the above statistics is not optimal, one might consider checking mapping and filtering parameters or further optimizing the Micro-C experiment. The summary of Micro-C interactions in this manuscript is available in **Supplemental Table 2**.

Valid Micro-C contacts were assigned to the corresponding ‘pseudo’ nucleosome bin. The bin file was pre-generated from the mouse mm10 genome by a 100-bp window that virtually resembles the nucleosome resolution. The binned matrix can be stored in HDF5 format as a COOL file by using the COOLER package (https://github.com/mirnylab/cooler)15 or in HIC file format by using the JUICER package (https://github.com/aidenlab/juicer)16. Contact matrices were then normalized by using iterative correction (IC) in COOL files^17^ or Knight-Ruiz (KR) in HIC files^18^. Regions with low mappability and high noise were blocked before matrix normalization. We expect that matrix balancing normalization corrects systematic biases such as nucleosome occupancy, sequence uniqueness, GC content, or crosslinking effects^17^. We notice that both normalization methods produce qualitatively equal contact maps. To visualize the contact matrices, we generated a compilation of COOL files with multiple resolutions (100-bp to 12,800-bp bins) that can be browsed on the Higlass 3D genome server (http://higlass.io)19. In this study, all snapshots of Micro-C or Hi-C contact maps and the 1D browser tracks (e.g., ChIP-seq) were generated by the HiGlass browser unless otherwise mentioned.

We evaluated the reproducibility and data quality for the Micro-C replicates using two published methods independently (https://github.com/kundajelab/3DChromatin_ReplicateQC)20. In brief, QuASAR-QC calculates the correlation of values in two distance-based transformed matrices. GenomeDISCO measures the difference in two graph diffusion-smoothed contact maps. We computed the matrix similarity scores between the biological replicates or between the untreated and IAA-treated cells for the 10-kb, 25-kb, 50-kb, and 250-kb Micro-C data. The detailed descriptions can be found in Sauria et al. for QuASAR-QC^21^ and Ursu et al. for GenomeDISCO^22^.

To analyze the genome-wide contact decaying *P*(s) curve, we used intra-chromosomal contact pairs to calculate the contact probability in bins with exponentially increasing widths from 100 bp to 100 Mb. Contacts shorter than 100 bp were removed from the analysis in order to minimize noise introduced by self-ligation or undigested DNA products. The orientations of ligated DNA are parsed into ‘IN-IN (+/-),’ ‘IN-OUT’ (+/+),’ ‘OUT-IN’ (-/-),’ and ‘OUT-OUT’ (−/+)’ according to the readouts of Illumina sequencing^23,24^. ‘UNI’ pairs combine ‘IN-OUT’ and ‘OUT-IN’ because both orientations are theoretically interchangeable. In this study, we plotted the contact decaying curves with the ‘UNI’ pairs and then normalized to the total number of valid contact pairs. Slopes of contact decay curves were obtained by measuring slopes in a fixed-width window searching across the entire range of decaying curves. We then plotted the derivative slope in each window against the corresponding genomic distance.

To identify chromosome compartments, we first transformed the Observed/Expected Micro-C matrices at the 200-kb resolution to the Pearson’s correlation matrices, and then obtained the eigenvector of the first principal component of the Pearson’s matrix by Principal Component Analysis (PCA). The sign of the eigenvector was corrected using active histone marks (H3K27ac and H3K4me3), as positive values are the A compartment (gene-rich or active chromatin) and negative values are the B compartment (gene-poor or inactive chromatin). The detailed description can be found in Lieberman-Aiden et al.^25^ The genome-wide compartment strength analysis shown as a saddle plot represents the rearrangement and aggregation of the genome-wide distance-normalized contact matrix with the order of increasing eigenvector values. The chromosome arm is first divided into quantiles based on the compartment score. All combinations of quantile bins are averaged and rearranged in the saddle plot. The Cooltools package (https://github.com/mirnylab/cooltools) has implemented the ‘call-compartments’ and ‘compute-saddle’ functions with the COOL files.

To identify chromatin domains (TADs) along the diagonal, we used insulation score analysis from the Cooltools package (https://github.com/mirnylab/cooltools) or arrowhead transformation analysis from the JUICER package (https://github.com/aidenlab/juicer)16. The detailed methods were described in Crane et al. for the insulation score analysis^26^ or Rao et al. for the arrowhead transformation analysis^27^. Briefly, we analyzed the insulation profile by using a 1-Mb sliding window that scans across Micro-C contact matrices at 20-kb resolution and assigns an insulation intensity to its corresponding bin. The insulation scores were obtained and normalized as the log2 ratio of the individual score to the mean of the genome-wide averaged insulation score. Chromatin boundaries can be identified by finding the local minima along with the normalized insulation score. Boundaries overlapping with low mappability regions were removed from the downstream analysis. The arrowhead analysis defines A*_i,i+d_* = (M\**_i,i-d_*–M\**_i,i+d_*)/(M\**_i,i-d_*+ M\**_i,i+d_*), where M* is the normalized contact matrix. A*_i,i+d_* can be thought of as the measurement of the directionality preference of locus *i*, restricted to contacts at a linear distance of *d*. A*_i,i+d_* will be strongly positive/negative if either *i,i-d* or *i,i+d* is inside the domain and the other is not, but A*_i,i+d_* will be close to zero if both loci are inside or outside the domain. By assigning this query across the genome, the edges of a domain will be sharpened and TADs can be detected. For aggregate domain analysis (ADA), each domain was rescaled to a pseudo-size by *N_i,j_*=((*C_i_*-*D_start_*)/(*D_end_*-*D_start_*), (*C_j_*-*D_start_*)/(*D_end_*-*D_start_*)), where *C*_i*,j*_ is a pair of contact loci within domain *D* that is flanked by *D_start_* and *D_end_*, and *N_i,j_* is a pair of the rescaled coordinates. The rescaled domains can be aggregated at the center of the plot with ICE or distance normalization. Coolpup (https://github.com/open2c/coolpuppy)28 has implemented a handy function to perform ADA with the COOL file. The lists of TAD called by the insulation score analysis or Arrowhead are available in **Supplemental Table 3 or 4**.

To identify loops/dots, we tested two novel algorithms, Mustache (https://github.com/ay-lab/mustache)29 and Chromosight (https://github.com/koszullab/chromosight)30, for the high-resolution Micro-C data. We found that both approaches outperform the HICCUPS algorithm in the JUICER package^16^ and the ‘Call-dots’ function in the Cooltools package in sensitivity and specificity to discover focal contact enrichment. In Mustache analysis, we called loops with balanced contact matrices at resolutions of 400 bp, 600 bp, 800 bp, 1 kb, 2 kb, 4 kb, 10 kb, and 20 kb using the calling options `--pThreshold 0.1 -– sparsityThreshold 0.88 -–octaves 2`. We then combined all loops at different resolutions. If an interaction was detected as a loop at different resolutions, we retained the precise coordinates in finer resolutions and discarded the coarser resolution. In Chromosight analysis, we used the ‘detect’ function to call loops with balanced contact matrices at resolutions of 400 bp, 600 bp, 800 bp, 1 kb, 2 kb, 4 kb, 10 kb, and 20 kb using calling options listed in **Supplemental Table 5**. We then combined all loops at different resolutions by the same approach as described above. We applied the lists of loop anchor to many downstream analyses by using Bedtools^31^, R, Python, or MATLAB, including (1) comparison of loop anchors between Micro-C and Hi-C or between different chromatin states; (2) distribution of loop strength or length; (3) cross-correlation with ChIP-seq, RNA-seq, and mNET-seq data; (4) ratio of boundary crossing, etc. For analysis of paired genomic loci (e.g., paired ChIP-seq peaks, genetic features, etc.) within a distance ranging from 2 kb to 2 Mb, we used Chromosight’s ‘quantify’ function to measure the probability of loop pattern for all intersections quantitatively. The loops were filtered by the following parameters: loop score >0.35 for 10-kb resolution, loop score >0.3 for 4-kb resolution, loop score >0.2 for 2-kb resolution, and the q-value lower than 10^−5^ for all resolutions (**Supplemental Table 5**). For aggregate peak analysis (APA) to assess genome-wide loop intensity, loops were centered and piled up on a 20-kb x 20-kb matrix with 400-bp resolution balanced data or 50-kb x 50-kb matrix with 1-kb resolution balanced data. Contacts close to the diagonal were excluded and normalized by a random shift matrix to avoid distance decay effects. The ratio of loop enrichment was calculated by dividing normalized center contacts in a searching window by the normalized corner submatrices. We used the same approach and normalization method to analyze the genome-wide target-centered loop intensity. Instead of aggregating at the intersection of loop anchors, the matrix is centered at the paired ChIP-seq peaks or genomic features. Coolpup (https://github.com/open2c/coolpuppy)28 has implemented the APA function for the COOL file. The lists of loops called by Mustache or Chromosight are available in **Supplemental Table 6 or 7**. The lists of loop quantification for cohesin, E-P, and P-P loops are available in **Supplemental Table 8 – 10.**

### Definition of chromatin states and structure observed by Micro-C

We first used the published ChromHMM (http://compbio.mit.edu/ChromHMM)32,33 to define the chromatin states in mESCs, which subclassifies chromatin into 12 states including: 1) CTCF/insulator, 2) active promoter (designated as “P”), 3) strong enhancer, 4) medium enhancer, 5) weak enhancer, 6) mix of promoter and enhancer, 7) bivalent promoter, 8) gene body, 9) Polycomb repressor, 10) intergenic regions, 11) heterochromatin, and 12) repeats. To simplify the analysis, we further combined the groups of strong, medium, and weak enhancers and mix of promoter and enhancer into “enhancer” (designated as “E”). In this study, we use the terms that are widely accepted in the field to describe the chromatin structures in Micro-C contact maps as well as avoid any ambiguous description that implicates their biological functions if they have not been well characterized, including: 1) Topologically-associating domain (TAD): squares along matrix diagonal enriched with self-interactions, which are defined as genomic intervals demarcated by the boundaries characterized by the insulation score analysis or the arrowhead transformation analysis; 2) Cohesin loops: focal enrichment of contacts in contact maps with the co-enrichment of CTCF/cohesin ChIP-seq peaks at loop anchors, which is thought to be formed by active loop extrusion halted by CTCF; 3) E-P/P-P loops: focal enrichment of contacts in contact maps with the co-enrichment of chromatin states for “active promoter (P)” or “enhancer (E)” at loop anchors. Although not all cohesin loops and E-P/P-P loops are formed through “looping” and some studies suggest using “dots” instead of “loops”, to simplify and be consistent with the majority of findings, we chose to use “loops” over “dots” to describe these enhanced focal contacts in this manuscript.

### RNA-seq experiments and analysis

Total RNA was extracted from ∼1×10^7^ mES cells (∼70% confluent P10 dish) with the standard TRIzol RNA extraction protocol. The abundant rRNAs were depleted from the sample using NEBNext rRNA Depletion Kit (NEB, #E6310). The rRNA-depleted RNAs were then subjected to RNA-seq library construction using the NEBNext® Ultra II Directional RNA Library Prep Kit for Illumina® (NEB, #E7765). The final RNA-seq libraries were amplified with 7 – 8 PCR cycles.

For RNA-seq analysis, we used Kallisto^34^ to quantify the number of transcripts and performed Sleuth^35^ analysis for DEGs identification according to the recommended settings in the walkthrough (https://pachterlab.github.io/sleuth_walkthroughs/). The DEGs were identified using the Wald test with the *q*-value < 0.01.

### Nascent RNA-seq experiment and analysis

We used the nascent RNA-seq (mNET-seq) protocol described in Nojima et al^36^ with minor changes. In brief, the chromatin fraction was purified from 1×10^7^ mES cells by the following procedure: 1) Wash cells with cold PBS twice; 2) Resuspend cells with 4 mL cold HLB+N (10mM Tris-HCl (pH 7.5), 10 mM NaCl, 2.5 mM MgCl_2_ and 0.5 % NP-40) and incubate for 5 min on ice; 3) Add 1 ml cold HLB+NS (10 mM Tris-HCl (pH 7.5), 10 mM NaCl, 2.5 mM MgCl_2_, 0.5 % NP-40 and 10 % Sucrose) under the layer of cell lysate; 4) Spin down cells and collect the nuclear pellet; 5) Add 120 µL cold NUN1 (20 mM Tris-HCl (pH 7.9), 75 mM NaCl, 0.5 mM EDTA and 50 % Glycerol) and transfer sample to a new tube; 6) Add 1.2 mL cold NUN2 (20 mM HEPES-KOH (pH 7.6), 300 mM NaCl, 0.2 mM EDTA, 7.5 mM MgCl_2_, 1 % NP-40 and 1 M Urea) and incubate for 15 min on ice while vortexing every 3 min; 7) Spin down the chromatin pellet and wash with cold PBS once. Next, chromatin and RNA were digested in 100 µL MNase digestion solution (1x micrococcal nuclease (MNase) buffer and 40 units/µl MNase (NEB, #M0247)) for 5 min at 37 °C while shaking at 1,400 rpm in a thermomixer. 25 mM EGTA was then added to inactivate the reaction. Digested/solubilized chromatin was then collected by centrifugation at 16,000xg for 5 min at 4 °C. The chromatin bound-Pol II complex was purified by the following steps: 1) Dilute 100 µL of sample with 400 µL cold NET-2 (50 mM Tris-HCl (pH 7.4), 150 mM NaCl, 0.05 % NP-40 and 1% Empigen BB (Sigma, cat no. 30326)); 2) Add 100 µL Pol II antibody-conjugated beads (10 µg Pol II 8WG16 antibody conjugated with Dynabeads™ Protein G for Immunoprecipitation (ThermoFisher, #1004D)) and incubate on a tube rotator for 1 hour in the 4 °C room; 3) Wash beads with 1 mL cold NET-2 for a total of 6 washes. RNA was then phosphorylated with T4 polynucleotide kinase (T4 PNK) with the following steps: 1) Wash beads with 50 µL cold 1X PNKT buffer (1X T4 PNK buffer and 0.1 % Tween); 2) Resuspend beads with 50 µL PNK reaction mixture (1X T4 PNKT, 1 unit/µL PNK and 1 mM ATP) for 5 min at 37 °C while shaking at 1,200 rpm in a thermomixer; 3) Wash beads with 1 mL cold NET-2. RNA was isolated by the standard TRIzol RNA extraction protocol with isopropanol RNA precipitation. Purified RNA was then dissolved in 10 µL of Urea dye (7 M Urea, 0.05 % Xylene cyanol, 0.05 % Bromophenol blue) and resolved on a 6% TBE-Urea gel at 200 V for 5 min. To size select 30-160 nt RNAs, we cut the gel between the Bromophenol blue and the Xylene cyanol dye markers. A 0.5-mL tube was punctured with 3-4 small holes by 26G needle and inserted in a 1.5-mL tube. Gel fragments were placed in the 0.5-mL tube and broken down by centrifugation at 16,000xg for 1 min. RNA was eluted by RNA elution buffer (1 M NaOAc and 1 mM EDTA) for 1 hr at 25°C in a Thermomixer shaking at 900 rpm. Eluted RNA was purified with SpinX column (Coster, #8160) with a glass filter (Whatman, #1823-010). The eluted RNA was purified again with ethanol precipitation. RNA libraries were prepared according to the protocol of the NEBNext Small RNA Library Prep Kit (NEB, #E7330). The mNET-seq library was obtained by PCR for 12 – 14 cycles.

For mNET-seq analysis, we wrote a custom pipeline to process raw data as follows: 1) Adapter trimming: we used TrimGalore (https://github.com/FelixKrueger/TrimGalore) to remove sequencing adapters ‘AGATCGGAAGAGCACACGTCTGAACTCCAGTCAC’ and ‘GATCGTCGGACTGTAGAACTCTGAAC’ at each side of the reads; 2) Mapping: Trimmed reads were mapped to the mouse mm10 reference genome with STAR RNA-seq aligner^37^; 3) Identifying the last nucleotide incorporated by Pol II: We used the Python script mNET_snr (https://github.com/tomasgomes/mNET_snr) to locate the 3’ nucleotide of the second read and the strand sign of the first read. The bigWig files were generated by using Deeptools as described above. To identified DEGs in mNET-seq, we used the NRSA (Nascent RNA Sequencing Analysis)^38^ package to statistically quantify differential changes of mNET-seq signal at the gene body between UT- and IAA-treated cells.

### Single-particle imaging experiments

All single-molecule imaging experiments were performed with a similar setting as described in our previous studies^4,39^. In brief, the experiments were performed with a custom-built Nikon TI microscope equipped with a 100X/NA 1.49 oil-immersion TIRF objective (Nikon apochromat CFI Apo TIRF 100X Oil), an EMCCD camera (Andor iXon Ultra897), a perfect focus system to correct for axial drift and motorized laser illumination (Ti-TIRF, Nikon), and an incubation chamber maintaining a humidified 37 °C atmosphere with 5% CO_2_ for the sample and the objective. Excitation was achieved using the following laser lines: 561 nm (1 W, Genesis Coherent) for JF549/PA-JF549 and TMR dyes; 633 nm (1 W, Genesis Coherent) for JF646/PA-JF646 dyes; 405 nm (140 mW, OBIS, Coherent) for all photo-activation experiments. Laser intensities were controlled by an acousto-optic Tunable Filter (AA Opto-Electronic, AOTFnC-VIS-TN) and triggered with the camera TTL exposure output signal. Lasers were directed to the microscope by an optical fiber, reflected using a multi-band dichroic (405 nm/488 nm/561 nm/633 nm quad-band, Semrock) and focused in the back focal plane of the objective. Emission light was filtered using single band-pass filters placed in front of the camera (Semrock 593/40 nm for TMR and JF549/PA-JF549 and Semrock 676/37 nm for JF646/PA-JF646). The angle of incident laser was adjusted for highly inclined laminated optical sheet (HiLo) conditions^40^. The microscope, cameras, and hardware were controlled through the NIS-Elements software (Nikon).

For ‘fast-tracking’ stroboscopic illumination (spaSPT) at ∼133 Hz, the excitation laser (633 nm for PA-JF646 or 561 nm for PA-JF549) was pulsed for 1 ms at maximum (1 W) power at the beginning of the frame interval, while the photoactivation laser (405 nm) was pulsed during the ∼447 µs camera transition time. Each frame consisted of a 7-ms camera exposure time followed by a ∼447-µs camera inactive time. The camera was set for frame transfer mode and vertical shift speed at 0.9 µs. With this setup, the pixel size after magnification is 160 nm and the photon-to-grayscale gain is 109. Typically, 30000 frames with this sequence were collected per nucleus, during which the 405-nm intensity was manually tuned to maintain an average molecule density of ∼0.5 localizations per frame, corresponding to ∼15,000 localizations per cell per movie. Maintaining a very low density of molecules is necessary to avoid tracking errors.

For ‘slow-tracking’ (slow-SPT) experiments, long exposure times (50 ms, 100 ms, and 250 ms) and low constant illumination laser intensities (0.5% - 2% of 0.5 W power) were used to measure residence time. The camera was set for normal mode and vertical shift speed at 3.3 µs. We generally recorded each cell with 1200 frames for a 250 ms exposure time, 3000 frames for a 100 ms exposure time, or 6000 frames for a 50 ms exposure time. We included H2B-HaloTag cells for the photobleaching correction for each experiment.

For PALM experiments, continuous illumination was used for both the main excitation laser (633 nm for PA-JF646 or 561 nm for PA-JF549) and the photo-activation laser (405 nm). The intensity of the 405 nm laser was gradually increased over the course of the illumination sequence to image all molecules and avoid too many molecules being activated at any given frame. The camera was set for 25-ms exposure time, frame transfer mode, and vertical shift speed at 0.9 µs. In total, 40,000–60,000 frames were recorded for each cell (∼20–25 min), which was sufficient to image and bleach all labeled molecules.

### spaSPT analysis

For analysis of spaSPT experiments, we used the QUOT package (https://github.com/alecheckert/quot) to generate trajectories from raw spaSPT movies with the steps of spot detection, subpixel localization, and tracking. All localization and tracking for this manuscript were performed with the following settings: 1) Detection: generalized log likelihood ratio test with a 2D Gaussian kernel (‘`llr`’ with k = 1.0, pixel window size (w) = 15, and a log ratio threshold (t) = 26.0); 2) Subpixel localization: Levenberg-Marquardt fitting of a 2D integrated Gaussian point spread function model (‘`ls_int_gaussian`’ with pixel window size (w) = 9, sigma = 1.0, ridge = 0.001, maximal iterations = 20 per PSF, and damping term = 0.3). 3) Tracking: we chose to use a conservative tracking algorithm with a 1.3-µm search radius (‘`conservative`’ with search radius = 1.3 and maximal blinks = 0). This setting makes the algorithm search for spot reconnections unambiguously, meaning that no other reconnections are possible within the specified search radius. Jumps are discarded if other reconnection possibilities given the search radius exist.

We next used the Spagl package (https://github.com/alecheckert/spagl)41 to estimate the likelihood of diffusion coefficients for each trajectory. The detailed discussion is available in Heckert et al^41^. In brief, we applied “State Array (SA)”, a grid of state parameters that span a range of diffusion coefficients (0 to 100 µm^2^/s), to calculate the posterior occupations of each point in the grid. The SA method conceptually produces a similar result as the Dirichlet process mixture models (DPMM) and retains its ability to model complex diffusive mixtures, while mitigating the issue of expensive likelihood functions. Instead of allowing infinite number of states (*K*→∞), the method fixes the number of states at a large but finite value. For each state *j* = 1, …, *K*, the algorithm chooses a fixed set of state parameters *θ_j_*. The model simplifies to:

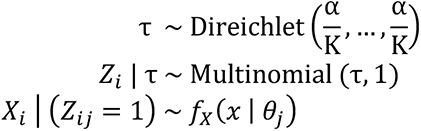

As a result, the SA method can compute more complex likelihood functions that incorporate localization error. Next, the algorithm infers the posterior *p* (Z, τ | X) with the Dirichlet prior *p*(τ) and corrects the systematic overestimation for the occupations of slow states due to defocalization. By marginalizing the posterior distribution on localization error, the method naturally incorporates uncertainty about the localization error of different states.

Alternatively, we analyzed the spaSPT data with the kinetic modeling framework implemented in the Spot-On package^39^. Briefly, the model infers the diffusion constant and relative fractions of two or three subpopulations from the distribution of displacements computed at increasing lag times (1Δτ, 2Δτ,…). This is performed by fitting a semi-analytical model to the empirical histogram of displacements using non-linear least squares fitting. Defocalization is explicitly accounted for by modeling the fraction of particles that remain in focus over time as a function of their diffusion constant. We used the following setting for Spot-On analysis in the manuscript: TimePoints = 8; BinWidth = 0.010; JumpsToConsider = 4; MaxJump = 5.05; ModelFit = CDF-fitting; NumberOfStates = 3; FitIterations = 3; FitLocErrorRange = 0.010-0.075; LocError = 0.035; Dbound range = 0.0001-0.05; Dfree range =0.5-25.

### Slow-SPT analysis

For analysis of slow-SPT experiments, we used the following tracking settings for this manuscript: 1) Detection: ‘`llr’` with k = 1.0, w = 15, t = 18; 2) Subpixel localization: ‘`ls_int_gaussian`’ with w = 9, sigma = 1.0, ridge = 0.001, maximal iteration = 20, and damp = 0.3; 3) Tracking: ‘`euclidean’` with search radius = 0.5, maximal blinks = 1, and maximal diffusion constant (µm^2^/s) = 0.08.

To extract residence times from slow-SPT data, we used long exposure times (50 ms, 100 ms, or 250 ms) to motion-blur freely diffusing molecules into the background^4,42–44^. We then recorded the trajectory length of each ‘bound’ molecule and used these to generate a survival curve (1-CDF), and performed double-exponential fitting to estimate the unbinding rates for non-specific binding (*K_ns_*) and specific binding (*K_s_*). We note that localization errors can cause both false-positive and false-negative detections. The *K_ns_* is likely to be contaminated by localization errors (e.g., from molecules close to being out-of-focus) and experimental noise. To filter out contributions from tracking errors and slow-diffusing molecules, we applied an objective threshold as previously described to consider only particles tracked for at least *N_min_* frames. To determine *N_min_*, we plotted the inferred residence time as a function of *N_min_* and observed convergence to a single value after ∼2.5 s. We thus used this threshold to determine the value of *K_s_*. To correct the biases from photobleaching, cell drifting, and background fluctuating, we assume that all these factors should affect H2B-HaloTag to the same extent as those affecting YY1-HaloTag. We can use an apparent unbinding rate for H2B-HaloTag as *K_bias_*, consistent with our FRAP analysis. Thus, we performed the slow-SPT experiments for YY1-HaloTag and H2B-HaloTag with the same camera and laser settings on the same day. We then obtained the residence time as:

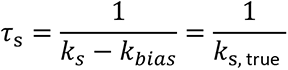

### Fluorescence recovery after photobleaching (FRAP) imaging analysis

FRAP was performed on an inverted Zeiss LSM 900 Axio Observer confocal microscope equipped with Airyscan 2 detector, a motorized stage, a full incubation chamber maintaining 37°C/5% CO_2_, a heated stage, an X-Cite 120 illumination source as well various laser lines. Images were acquired on a 40x Plan NeoFluar NA1.3 oil-immersion objective at a zoom corresponding to a 76 nm x 76 nm pixel size. The microscope was controlled using the Zeiss Zen imaging software.

In this manuscript, we recorded 60 sec of movies for YY1-HaloTag at one frame per 250 ms, corresponding to a total of 240 frames. The first 20 frames were acquired before the bleach pulse, allowing us to accurately measure baseline fluorescence. A circular bleach spot (r = 6 pixels) was chosen in a region of homogenous fluorescence at a position at least 1 µm from nuclear or nucleolar boundaries. Alternatively, we bleached a square at one corner of nucleus, which reduces noise while introducing some uncertainty for our downstream fitting analysis. The spot was bleached using maximal laser intensity and pixel dwell time corresponding to a total bleach time of ∼1 s. We note that because the bleach duration was relatively long compared to the timescale of molecular diffusion, it is not possible to accurately estimate the bound and free fractions from our FRAP curves.

To quantify FRAP movies, we wrote a pipeline in MATLAB. Briefly, our algorithm automatically detects the bleached spot, the background spot, and the nucleus segments by Gaussian smoothing, hole-filling, and segmenting a nucleus in a FRAP movie. Cell drift is also automatically corrected by the optimal linear translation in x and y. Next, we quantify the bleach spot signal as the mean intensity of a slightly smaller region, which is more robust to lateral drift. The FRAP signal is corrected for photobleaching using the measured reduction in total nuclear fluorescence and internally normalized to its mean value during the 20 frames before bleaching. Finally, corrected FRAP curves from each single cell were averaged to generate a mean FRAP recovery. We used the mean FRAP recovery in all figures and for model-fitting.

Model selection is critical to infer the parameters from FRAP experiments. Sprague et al.^45^ suggested that when:

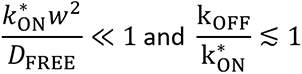

Then a ‘reaction dominant’ FRAP model is most appropriate. For YY1:

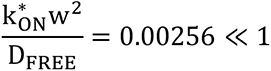

Thus, a reaction-dominant FRAP model is the most suitable choice for YY1’s FRAP modeling. Sprague et al.^45^ demonstrated that the FRAP recovery depends only on *k_OFF_* in the reaction-dominant regime. We thus fit the FRAP curves to the model and applied the slower off rate to estimate the residence time according to 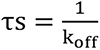.

### Inferring parameters related to YY1’s target search mechanism

We used the parameters inferred from our spaSPT and the residence time measurements from our FRAP or slow-SPT analysis. The detailed discussion is available in Hansen et al^4^. Briefly, from the Spagl State Array analysis, we determined that the total bound fraction for YY1 is ∼31%. However, the total bound fraction (0 – 0.1 µm^2^/s) contains both YY1 molecules bound specifically to their target sites and non-specific interactions (e.g., sliding or jumping on DNA). We previously estimated the fraction that is non-specifically bound using a mutant CTCF with a His-to-Arg mutation in each of the 11 zinc-finger domains^4^. This CTCF mutant is virtually unable to interact specifically with its binding sites. The Spagl analysis estimated the bound fraction to be ∼8.1% for this mutant in mESCs. Since we did not perform the spaSPT experiments for YY1’s DNA binding domain mutants, we thus inferred the *F_BOUND,specifc_* ∼= *F_BOUND,total_* in this manuscript.

We next determined the average time for YY1 to find its cognate site after dissociating from the previous site. We will use ‘*s*’ and ‘*ns*’, as abbreviations for specific and non-specific binding, respectively, in the following discussion. The pseudo-first-order rate constant for specific binding sites, *k^∗^*_ON,s_, is related to the fraction bound by:

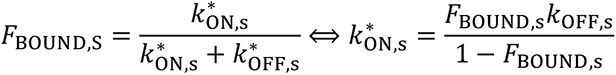

We determined the off-rate for a specific interaction in our residence time measurements. Thus, we can calculate *k^∗^*_ON,s_, which is directly related to the average search time for a specific YY1-binding site:

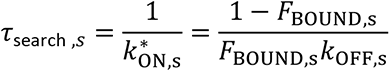

After plugging in these determined parameters of *F_BOUND,s_* and *k*_OFF,s_, we obtained total search times for YY1 of ∼28.3 s in wild type mES cells, ∼31.6 s in CTCF-depleted cells, ∼61.4 s in RAD21-depleted cells. We inferred the residence time estimated from the slow-SPT data with 100 ms exposure time or the FRAP analysis in this manuscript.

### PALM analysis

For analysis of PALM experiments, we used the publicly available ThunderSTORM package (https://github.com/zitmen/thunderstorm)46 with the following setting for this manuscript: 1) Image filttering: ‘`Wavelet filter (B-Spline)`’ with B-Spline order = 3 and B-Spline scale = 2.0; 2) Approximate localization: ‘`Local maximum`’ with peak intensity threshold = 1.5*std(Wave.F1) and 8-neighbourhood connectivity; 3) Subpixel localization: ‘`Integrated Gaussian`’ with fitting radius = 3 pixels, fitting method = maximum likelihood, initial sigma = 1.6, multi-emitter analysis is disabled; 4) Image reconstruction: ‘Averaged shifted histogram’. After tracking, we further filtered ambiguous emitters with the following setting: 1) Filtering: `frame > 100 & intensity > 100 & sigma < 220 & uncertainty_xy < 50`; 2) Merge: `Max distance = 10 & Max frame off = 1 & Max frames = 0`; 3). Remove duplicates is enabled. This setting combines the blinking molecules into one and removes the multiple localizations in a frame.

### Antibodies

See **Supplementary Table 1** for a complete list of the antibodies used in this study.

### Datasets and accession numbers

The Micro-C, ChIP-seq, nascent RNA-seq and total RNA-seq data generated in this publication have been deposited in NCBI’s Gene Expression Omnibus and are accessible through GEO Series accession number GSE178982. We also reanalyzed data that we previously generated in wild type mESCs (GSE130275)^13^. spaSPT raw data are accessible through DOI: <u>10.5281/zenodo.5035837</u>.

